# Reference-guided Genome Assembly of Long Non-coding RNA Transcripts Reveals Target Genes Associated With Crohn’s Disease

**DOI:** 10.1101/2025.09.30.674088

**Authors:** Meaghan M. Kennedy Ng, Sophie Silverstein, Nina C. Nishiyama, Caroline Beasley, Grace Lian, Benjamin Huan, Gwen Lau, David Weaver, Ayesh Awad, Matthew R. Schaner, Shehzad Z. Sheikh, Terrence S. Furey

**Affiliations:** Curriculum in Bioinformatics and Computational Biology, Department of Genetics, School of Medicine, The University of North Carolina at Chapel Hill, Chapel Hill, North Carolina, USA; Center for Gastrointestinal Biology and Disease, Division of Gastroenterology and Hepatology, Department of Medicine, School of Medicine, The University of North Carolina at Chapel Hill, Chapel Hill, North Carolina, USA; Department of Biology, College of Arts and Sciences, The University of North Carolina at Chapel Hill, Chapel Hill, North Carolina, USA

**Author notes:** Corresponding Authors: Shehzad Z. Sheikh M.D., Ph.D. 7314 Medical Biomolecular Research Building, 111 Mason Farm Road, Chapel Hill, North Carolina 27599,; Terrence S. Furey, Ph.D., 5046 Genetic Medicine Research Building, 120 Mason Farm Road, Chapel Hill, North Carolina 27599.

**Keywords:** long non-coding RNAs, Crohn’s disease, transcriptomics

## Abstract

Crohn’s disease (CD) is highly heterogeneous in presentation and progression with no cure. Molecular phenotyping has been used to elucidate cellular and tissue-based alterations to characterize drivers and effects of disease. One currently understudied class of functional molecules is long non-coding RNAs (lncRNAs). Studying the full lncRNA landscape in IBD is challenging due in part to an incomplete lncRNA annotation and a lack of their functional characterization in tissues of interest. We used a genome-guided alignment strategy to assemble predicted lncRNA transcripts using short RNA-sequencing data from colon tissue of adult patient samples. When combining our predicted lncRNAs with previous lncRNA annotations, we determined 98 that were differentially expressed, recapitulating many from previous IBD studies while also uncovering new ones. We built gene co-expression networks to cluster lncRNAs with functionally characterized protein-coding genes. Clusters containing differential lncRNAs were correlated to disease status and associated with pathways related to the humoral immune response, metabolism, and tissue regeneration. We uncovered multiple differential lncRNAs whose expression significantly correlated with nearby differential protein-coding genes that have also been differentially expressed in other IBD datasets, such as *PITX2*. We focused on a predicted lncRNA that is antisense to the PITX2-adjacent lncRNA *PANCR*, which we called *PANCR-AS1*, and provide multiple lines of evidence that support *PANCR-AS1* functioning as an enhancer of *PITX2* expression. Overall, we determined lncRNAs that are potential contributors to CD pathogenesis. We developed a robust pipeline for identifying lncRNAs in diseased and non-diseased tissue that are absent from reference annotations. We also outlined a framework to pinpoint significant disease-associated lncRNAs with potential functional activity related to their nearby protein-coding genes.

## Background

Crohn’s disease (CD), a primary subtype of inflammatory bowel disease (IBD), is a chronic inflammatory condition caused by an inappropriate immune response within a genetically susceptible host to enteric microbiota[1], [2], [3]. CD is heterogenous in its clinical presentation that includes a broad spectrum of disease behaviors including the development of strictures that cause bowel obstructions and penetrating disease that can lead to fistulas and intestinal perforations[2], [3]. There is no cure for CD, and there exists an urgent need to better understand the molecular contributors to disease phenotypes.

Long non-coding RNAs (lncRNAs), a class of transcripts that are >500 base pairs that are not translated into proteins, have been shown to have diverse roles in the regulation of gene expression. These include modifying chromatin accessibility either directly through binding to the DNA sequence or indirectly by the post-translational modification of epigenetic markers such as histone tails to alter chromatin compaction; they also have been shown to help recruit transcription factors to regulatory elements for the activation or repression of target genes[4], [5], [6].

Multiple lncRNAs have been shown to be differentially expressed in patients with IBD compared to those without disease (non-IBD)[7], [8], [9], [10], [11]. Specific lncRNAs have been linked with ameliorating or promoting inflammation (*LUCAT1*, *IFNG-AS1/NeST,* and *NAIL*)[12], [13], [14], [15] and regulating intestinal homeostasis (*HOXA11os*, *IRF1-AS1/CARINH*)[16], [17]. Consequently, *IRF1-AS1, IFNG-AS1,* and other lncRNAs have been associated with enhancer-like activity in mitigating inflammatory response pathways[14], [16], [18]. These enhancer lncRNAs (e-lncRNAs) produce multi-exonic transcripts that are correlated to the activity of their enhancer elements and the expression of nearby protein-coding genes[19], [20], [21]. Evolutionary conservation of splice sites and the correlation of splicing efficiency to nearby gene expression support the importance of e-lncRNA splicing to its cis-regulatory activities[20], [21], [22]. Genome-wide association studies (GWAS) of IBD and other autoimmune diseases have shown that ∼90% of disease-associated genetic variants lie predominantly in non-coding regions including in enhancers[23], [24]. These may include e-lncRNA associated enhancers whose activity may provide a better understanding of the consequences of those variants. Despite these efforts, the role of lncRNAs in IBD, like most complex diseases, remains understudied due to the inherent challenges in comprehensively annotating and functionally characterizing them. Most have low and even nearly undetectable expression levels compared with protein-coding genes. Many are only expressed in certain cells or tissues, at certain developmental timepoints, or in certain disease contexts or conditions[5]. With improvements in RNA-sequencing technologies, the number of annotated lncRNA genes has been steadily rising, with estimates of there being 17000 to >90000 lncRNAs in the human genome[4]. Data from RNA-seq experiments can be used to assemble novel lncRNAs not currently annotated in reference transcriptome collections such as GENCODE[25], [26]. For well-annotated species such as humans, genome reference-guided assembly approaches such as *StringTie* and *Scallop*, and other tools have been successful in both recovering known transcripts and discovering new ones[27], [28]. To focus on lncRNAs that may have functional potential, priority should be given to lncRNAs that are (i) spliced; (ii) contain evolutionarily conserved elements in key locations such as the promoter region, 5’ end of the transcript, and around splice sites; (iii) show gene synteny across multiple species; and (iv) have cell-type specific chromatin marks, such as H3K4 mono- or tri-methylation, near their transcription start sites (TSS)[29].

The goal of this study was to provide a comprehensive analysis of lncRNAs, including currently unannotated transcripts that are predicted to be lncRNAs, in colon tissue of patients with and without Crohn’s disease (CD). To do this, we first generated a lncRNA transcriptome using RNA-seq data from an adult cohort of 90 CD patients and 50 non-IBD controls. Differential analyses of lncRNA expression between CD and non-IBD individuals supported both previously identified differentially regulated lncRNAs in IBD, as well as many uncharacterized in the IBD including some of our predicted lncRNA genes. Gene co-expression analysis with protein coding genes provided evidence for the role of CD-associated lncRNAs in commonly altered processes in IBD, including immune response pathways, metabolic processes, and tissue regeneration. Correlation of expression with nearby protein-coding genes predicted potential gene regulatory targets of some lncRNAs.

We highlight one particularly interesting differentially expressed lncRNA from our predicted set, which we called (*PANCR-AS1)*. This lncRNA is antisense to and in the intron of the previously annotated PITX2-adjacent lncRNA *PANCR*. Expression of *PANCR-AS1* is strongly positively correlated to the expression of the adjacent transcription factor *PITX2* gene, which has been found to be differentially expressed in IBD[30], [31]. Multiple lines of evidence support PANCR-AS1 as a functional lncRNA, including regulatory element conservation across species and overall sequence similarity in syntenic lncRNAs, cell-type specific chromatin accessibility and other regulatory profiles, and potential interactions between the promoter of *PANCR-AS1* and *PITX2*. We hypothesize that this lncRNA could act as an e-lncRNA to regulate the expression of *PITX2*.

## Methods

### Patient Recruitment and Sample Collection

90 CD and 50 non-IBD adult patients were recruited at the UNC Multidisciplinary Inflammatory Bowel Diseases Center. This study was approved by the Human Research Ethics Committee at UNC-Chapel Hill and was carried out in accordance with the Institutional Review Board (approval numbers 10-0355 and 11-0359). Samples were collected and stored in RNA-later from macroscopically non-inflamed sections of colonic mucosa for patients undergoing colonoscopy or surgical resection. RNA was isolated using the Qiagen RNeasy Mini Kit, as previously described[32]. All reads were 50 bp long, paired-end, and stranded. Quality control was performed using fastqc (v0.11.9)[33] and multiqc (v1.11)[34]. All samples have a median transcript integrity score of >65.

### Unannotated lncRNA Prediction

Unannotated lncRNAs were predicted using all RNA-seq data as illustrated in Figure 1A. Adapters were removed and low-quality bases (Q<10) were trimmed using Cutadapt (v.2.9.0)[35]. Trimmed reads <20 base pairs (bps) were removed. Reads were aligned to the GRCh38 human reference genome with STAR (version 2.7.9a)[36]. Transcripts were assembled within each sample using Stringtie (version 2.1.5)[27] and the GENCODE v39 reference transcriptome to guide assembly. Transcript fragments with expression <1 transcript per million (TPM) were removed. A meta-assembly of transcripts across samples was generated using TACO (v.0.7.3)[37]. Assembled transcripts were categorized relative to the reference transcriptome using GffCompare (v0.10.4)[38], and fully intronic antisense transcripts (“i"), antisense exon overlapping transcripts (“x”), and intergenic transcripts (“u”) were retained. Transcripts <500bps were filtered using Gffread[38]. The transcript coding potential was predicted using CPPred (v1)[39], CNCI (v2)[40], and PLEK (v1.2)[41], and transcripts predicted as coding by any method were filtered out.

**Figure 1.**
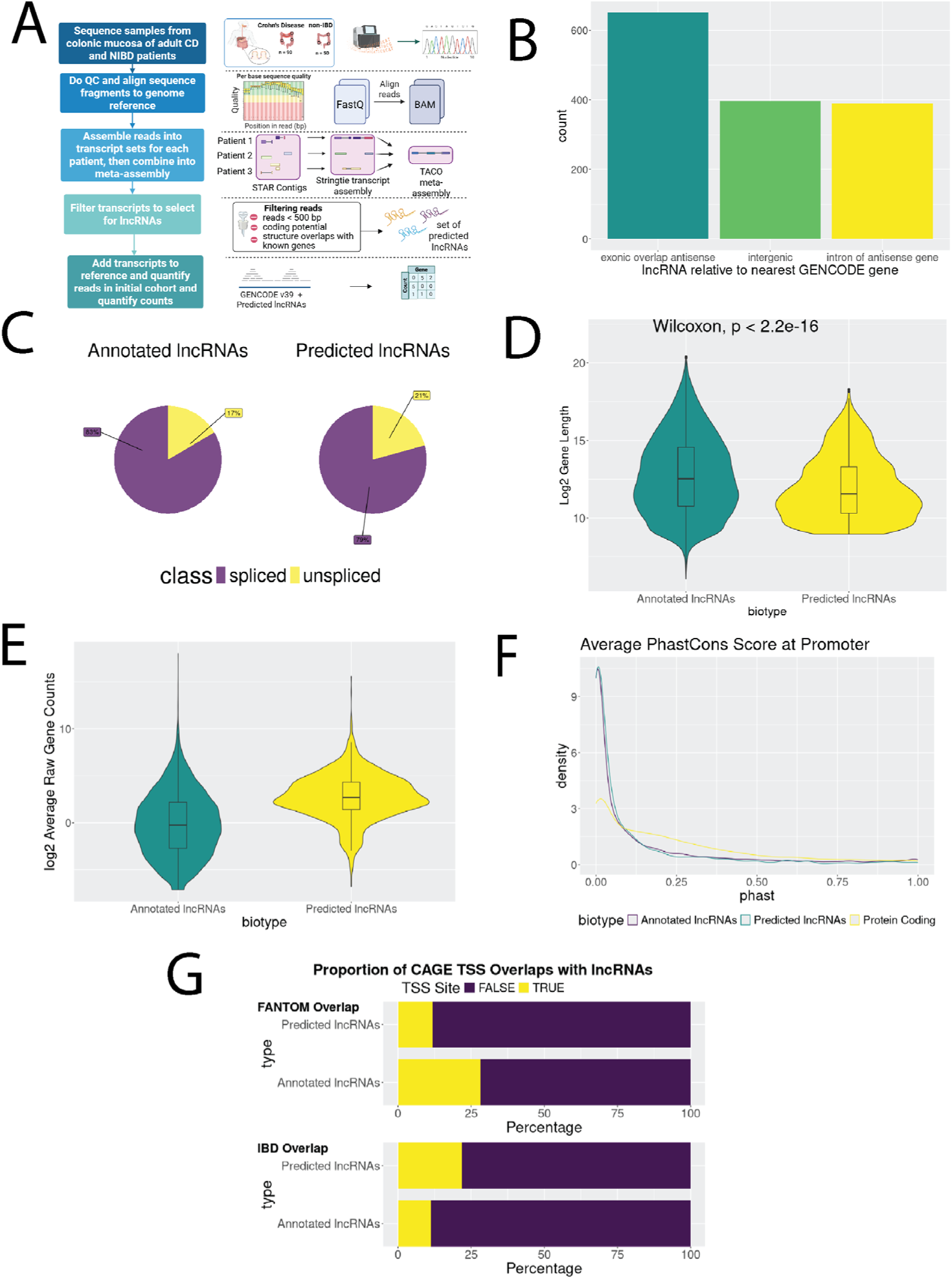
Comparisons of attributes of predicted lncRNAs with GENCODE v39 annotated lncRNAs. (A) Summary of the lncRNA discovery pipeline (created with BioRender) (B) Transcription orientations of predicted lncRNAs to GENCODE gene annotations. (B) Breakdown of lncRNA sub-categories (intergenic, intronic, antisense) by annotation. (C) Proportions of lncRNAs that are spliced or unspliced. Distribution of annotated and predicted lncRNAs based on their (D) transcript length and (E) mean TPM expression. (F) Proportion of lncRNA transcription start sites (TSS) with both FANTOM and IBD CAGE annotated TSS. (G) Comparison of vertebrate-level conservation scores of the promoter regions of predicted lncRNAs, GENCODE lncRNAs, and GENCODE protein-coding genes.

### Transcript Quantification

A decoy-aware index was built using a concatenated transcriptome of GENCODE v39 and the predicted lncRNAs using salmon index (Salmon v.1.6.0)[42], [43]. RNA-seq data was aligned and quantified using salmon quant in mapping-based mode. Estimated counts were imported into R (v4.3.1) using tximport (v1.26.1)[44], and lowly expressed genes, defined as those with a counts-per-million (CPM) of 70% of CD or non-IBD samples, were filtered out using *filterByExpr* (edgeR[45]).

### Annotating lncRNA candidates

Predicted and annotated (GENCODE v39) lncRNAs were annotated as follows:

1. <u>Cap analysis of gene expression (CAGE) data</u>: Coordinates of stringent transcript start sites (TSS) from FANTOM[46] and known and novel TSS determined by CAGE in IBD patients[47] were lifted from hg19 to hg38 using *liftOver* (from ucsctools, v320)[48]. Annotated TSS regions (+/-250 bp from the TSS) were determined using GenomicRanges (v1.52.1) and GenomicFeatures (v1.52.2)[49].
2. <u>Protein coding genes</u>: Pairs of significantly differentially expressed protein-coding genes and lncRNAs within 100kb of one another were determined using the *findOverlapPairs* function in the GenomicRanges[49] package. Pearson correlations between the estimated counts for the protein-coding and lncRNA gene pairs were determined using the *cor* function, calculations were made using the *cor.test* function, and adjusted p-values were calculated using the *p.adjust* function in the stats package in base R (v.4.3.1).
3. <u>Evolutionary conservation</u>: The average conservation in the region 200bp upstream of the TSS was determined using phastCons[50] scores from the 100-way species annotation (phastCons100way.UCSC.hg38 package in R)[51].
4. <u>Determining lncRNA subclasses:</u> Coordinates of annotated and predicted lncRNAs were intersected with annotated protein-coding genes using GffCompare[38]. Genes were determined to be intergenic if all transcripts were intergenic (class code “x”), and antisense if at least one transcript overlapped a protein-coding exon on the opposite strand (“u”). After retaining intronic (“i”) transcripts, the annotated protein-coding gene set was split into separate exon-only and transcript-only GTF files; bedtools[52] (v2.31.1) *subtract* was used to create a GTF file containing only the intronic regions, and bedtools *intersect* was used to determine whether the intronic regions were sense or antisense to the transcripts.

### Differential Expression Analysis

Sources of unwanted technical and biological variation were modeled and removed using RUVSeq (v1.34.0)[53]. Non-differential genes (nominal p-value of > 0.1) were used as control genes, which were determined using the likelihood ratio test in DESeq2 (v1.40.2)[54] and the full model = ∼sex + disease, reduced model ∼1. Differential analysis was performed using DESeq2 with a linear model that included sex and the first 4 RUV factors as covariates.

Independent hypothesis weighing was performed using IHW (v1.28.0)[55] to improve the power of multiple hypothesis testing. Genes with adjusted p-values < 0.05 were considered significant.

### Co-Expression Network Generation

Counts were normalized using the *varianceStabilizingTransformation* function from DESeq2, and covariates (including sex and the first 4 RUV factors) were regressed out using the *empiricalBayesLM* function in WGCNA (v1.72)[56]. The count matrix was filtered to remove the 25% least variable genes. The soft thresholding power was determined to be the power that had reached an R^2^ fit at 0.9 (Supplementary Figure 4A) from an R^2^ model fit while reducing the mean k connectivity to <100 (Supplementary Figure 4B). Network construction and module detection were performed using the *blockwiseModules* function (mergeCutHeight = 0.25, deepCut=4, power = 9, network type = "signed", corType = "bicor", maxPOutliers = 0.1).

A representation of the gene expression profile for each module (“eigengene”) were calculated using the *moduleEigengenes* function in WGCNA, and the correlation between module eigengenes and disease was calculated using the *bicor* function (use = "p", robustY = FALSE, maxPOutliers = 0.1) in WGCNA and plotted using the *labeledHeatmap* function in WGCNA. GO pathway analysis was done using the *enrichGO* function in the clusterProfiler (v4.8.3) package[57]. For IBD-associated clusters with > 20 significant terms, we summarized redundant terms using REVIGO[58]. Node and edge tables were exported using the *exportNetworkToCytoscape* function in WGCNA and were visualized in Cytoscape (v3.10.2)^49^.

### Conservation and Transcript Synteny

We analyzed the evolutionary conservation between the combined set of annotated and predicted lncRNAs and a set of mouse lncRNAs (GENCODE vM37, GRCm39 assembly) using the tool slncky[59]. In brief, *liftOver*[48] is used to find regions of synteny, or regions between protein-coding gene orthologs. If there are transcripts annotated in both species, the two syntenic regions are aligned (+/-150,000 bp) using *lastz* (v1.03.73)[60] with a reduced gap penalty to better tolerate insertions and deletions, and is compared with alignments of transcripts to shuffled intergenic regions as the null distribution (p < 0.05).

A BLASTN alignment between the PANCR-AS1 transcript sequence against the Ensembl non-coding gene databases in other species including i.) domestic cat ii.) goat iii.) horse and iv.) sheep was run using the BLAST/BLAT search tool in the Ensembl web browser (v114)[61], with the search sensitivity set for distant homologies.

Sequence conservation annotations of PANCR-AS1 regulatory regions were based on i.) the “Hiller Lab 470 Mammals Track”, which calculated conservation scores using phastCons[50], based on the alignments of 470 mammalian species using multiz[62]. The promoter region was further characterized into regions of high and low interest using a combination of the “combined annotation dependent depletion” (CADD) score[63] and the Zoonomia “runs of contiguous phyloP constraint” (RoCCs) score[64].

### Annotation of PANCR-AS1 Regulatory Activity Using Publicly-Available Data

ATAC-seq peaks across multiple human tissues, as well as precision run-on and capped RNA sequencing (PRO-cap) and chromatin interaction analysis by paired-end tag sequencing (ChIA-PET) data from the Caco-2 cell line, were sourced from the ENCODE repository[65], and ChromHMM[66], [67] calls were sourced from the Roadmap Epigenomics Project[68]; accession numbers are provided in Supplementary Table 1.

Significant ChIP-seq peaks (MACS2[69] adj. p < 1x10^-5^) for Pol II, NELFA, NELFCD, NELFE, SUPT5H, EZH2, and SUZ12 were downloaded from the ChIP-Atlas (v3.0) server[70].

To determine whether *PANCR-AS*1 interacts with PRC2 in tissues where repression is occurring, we looked at RNA immunoprecipitation sequencing (RIP-seq) data that captured RNA interactions with the PRC2 subunit EZH2 in mouse embryonic stem cells, which was sourced from Zhao et al.[71] Mouse coordinates were lifted from mm10 to mm39 using *liftOver*[48]. All publicly-available data were visualized using IGV[72], [73].

## Results

### Predicted lncRNAs show similar characteristics to annotated lncRNAs

To predict lncRNAs not currently annotated (GENCODE v39), potentially due to low expression levels and/or context specificity, we constructed a robust lncRNA discovery pipeline using RNA-seq data from 140 adult patient samples (Figure 1A, see Methods for details). Of the initial 78670 transcripts that were assembled, transcripts with complete matches to a reference gene annotation (27663, 35.2%) and partial matches to isoforms of an annotated gene (39659, 50.4%) were removed. The remaining 11348 transcripts (14%) were antisense, intronic, or intergenic relative to annotated genes (Supplementary Figure 1). Of these, transcripts that were <500 base pairs (bps) in length, identified as coding by at least one of three coding potential tools, and/or were intronic to an annotated gene but transcribed in the sense direction were filtered out, leaving a final set of 2520 predicted lncRNA transcripts representing 1436 unique genes.

We compared these with additional lncRNA annotated sets and found that most transcripts (2112, 83%) and genes (1237, 86%) were unannotated in at least one of the three references, while about a quarter of the transcripts (662, 26%) and genes (404, 28% genes) were unannotated in all three references (Supplementary Figure 2). This both supports the robustness of our transcripts while showing the relative incompleteness of annotated lncRNAs.

To further assess the quality of the predicted lncRNAs, they were quantified and compared with characteristics of 51932 annotated lncRNA transcripts (17755 genes; GENCODE v39; Figures 1B-G). Positionally, the majority of both transcript sets were intergenic in relation to protein-coding genes; these totaled 34227 out of 51932 (66.0%) of the annotated lncRNA transcripts, while making up 1371 out of 2520 (54.4%) of the predicted transcripts. Similar proportions of both transcript sets were antisense to and overlapped a protein-coding gene, totaling 10096 (19.4%) of the annotated and 385 (15.3%) of the predicted lncRNA transcripts. For the predicted transcript set, the majority of the rest were in introns of protein-coding genes, totaling 711 (28.2%) of the predicted set. While 5544 (10.7%) of annotated lncRNA transcripts were in the introns of protein-coding genes, the majority of the remaining annotated lncRNA transcript set belonged to other categories, including transcripts that appear to be non-coding isoforms of protein-coding genes, or long lncRNAs that contained protein-coding genes within an intron (Figure 1B). Lastly, 83% of annotated and 79% of predicted lncRNA genes showed evidence of splicing (Figure 1C).

We then analyzed transcript lengths of both lncRNA sets (Figure 1D) and found that while the annotated lncRNAs were longer on average (11568 bps predicted, 33805 bps annotated), both distributions were skewed by extremely long lncRNAs. Their median lengths were more similar (3022 bps predicted, 5858 bps annotated). Finally, we compared the mean expression patterns (Figure 1E). Mean normalized TPM expression was significantly higher in predicted lncRNAs (2.63) compared to annotated lncRNAs (1.20) when excluding genes with zero counts; this is expected because assembling transcripts from our cohort likely selected for colonic (tissue-specific) and disease-associated (context-specific) lncRNAs, whereas annotated lncRNAs are generated across numerous tissues and cell lines.

Finally, we compared the predicted lncRNAs to other publicly-available annotations. First, we looked at the sequence conservation in the promoter region of each lncRNA (+/-500 bp of the TSS) using the 100 vertebrate conservation scores from PhastCons[51]. We found that the distribution of promoter sequence conservation was similar in both annotated lncRNAs (median = 0.034, IQR = 0.002 - 0.183) and predicted lncRNAs (median = 0.026, IQR = 0.003 - 0.122) with lncRNAs being less conserved compared to protein-coding gene promoters (median = 0.169, IQR = 0.034 - 0.367) as expected (Figure 1F). Second, annotated and predicted lncRNA TSS were matched to annotations based on CAGE sequencing of tissues from the FANTOM consortium, as well as to a separate study that performed CAGE sequencing in colon tissue from IBD patients[47]. We found that while fewer annotated lncRNA TSS were supported by the IBD dataset than the FANTOM dataset (11.3% IBD vs 28.3% FANTOM), proportionally more predicted lncRNA TSS were supported in the IBD dataset (21.9% IBD vs 11.8% FANTOM; Figure 1G).

These analyses show that the predicted lncRNAs were similar to the annotated lncRNAs in terms of their transcript length, splicing activity, and sequence conservation patterns; however, there were noticeable increases in the overlap of our TSS sites to the IBD-specific CAGE sequencing dataset (as opposed to FANTOM), as well as an increase in the mean TPM counts of predicted lncRNAs as compared to annotated lncRNAs. Together, these provide support for the robustness of our discovery pipeline and annotations but also indicates a bias towards more context-specific lncRNAs in the predicted set.

### Annotated and predicted lncRNAs are differentially expressed in CD

We sought to identify lncRNAs that are differentially expressed in CD compared with non-IBD controls. We considered all genes, first filtering lowly expressed genes leaving 14635 protein-coding genes, 2120 previously annotated lncRNA genes and 537 of our predicted lncRNAs. We found 791 genes that were significantly differential (padj < 0.05; Figure 2A; Supplementary Table 3), of which 98 were lncRNAs (Figure 2B). Similarly, after correcting for unwanted sources of variation, we performed principal component analysis (PCA) on the 250 most variable lncRNAs and saw a partial separation between non-IBD and CD patients (Figure 2C).

**Figure 2.**
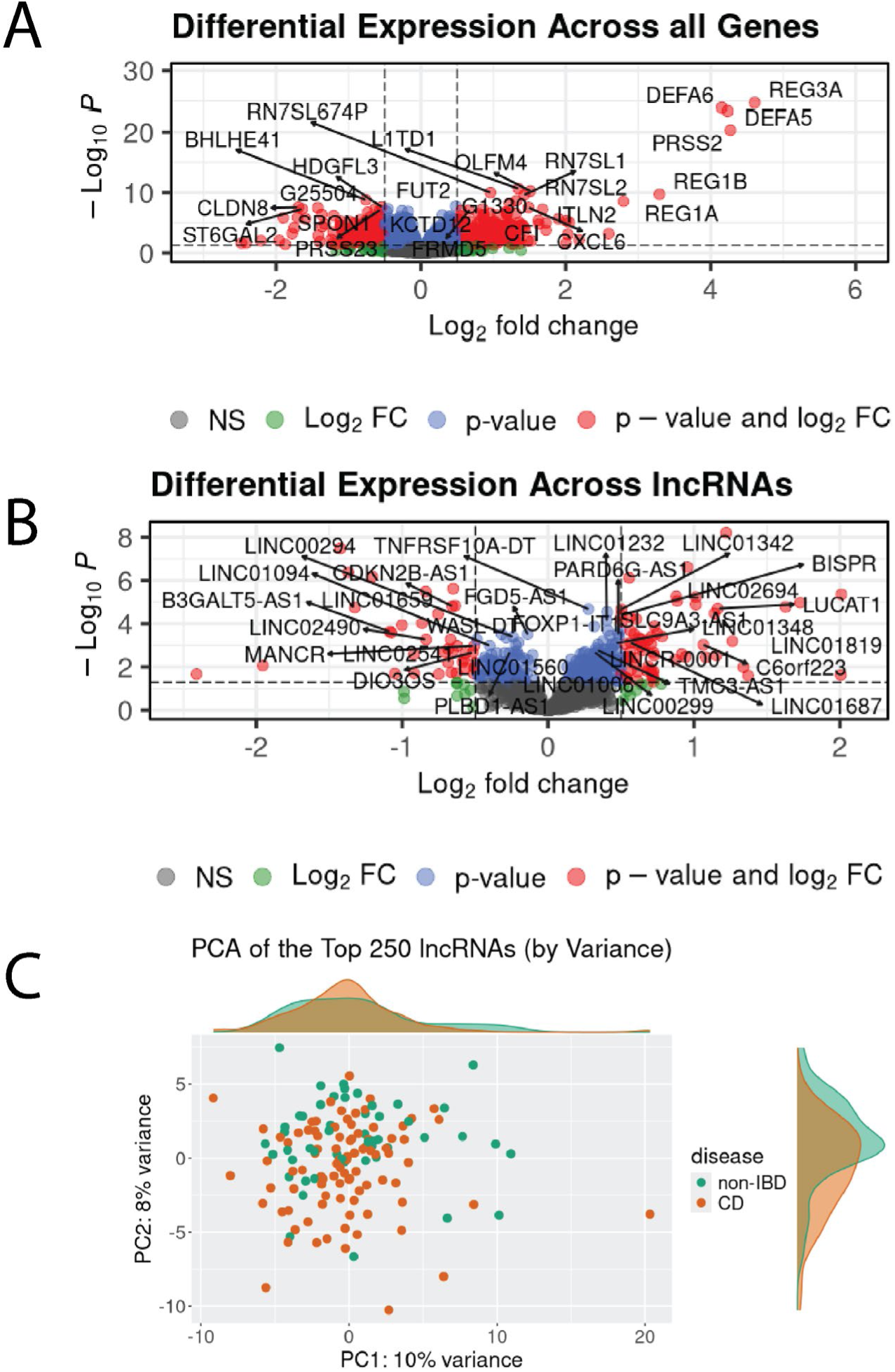
CD and non-IBD patients show group-wide changes in gene expression. (A) Volcano plot of all genes tested for differential expression in CD vs non-IBD. The top 25 most differential genes based on DESeq2 adjusted p-values are labelled. (B) Volcano plot of all lncRNA genes tested for differential expression in CD vs non-IBD. The top 25 genes based on DESeq2 adjusted p-values with an official gene name are labelled. (C) PCA of the top 250 most variably expressed lncRNA genes across all samples.

Of the 98 differential lncRNAs, 64 had increased expression in CD patients with a log2 fold change greater than zero (log2FC > 0), while 34 had decreased expression (log2FC < 0). Interestingly, 38 (39%) were from the predicted set. Of the differential annotated lncRNAs in our cohort, we found several that had been previously reported as differential and implicated in IBD, including *CDKN2B-AS1* (*ANRIL*), *DIO3OS*, and *LUCAT1*[11], [12]. However, the majority of lncRNAs have not been previously associated with IBD or immune-related diseases.

### Correlated expression suggests regulatory targets for lncRNAs

Many lncRNAs are known to regulate the expression of nearby genes. Therefore, we wanted to identify adjacent (<100Kb) differential protein-coding genes and lncRNAs that showed strong positively or negatively correlated expression indicating a possible regulatory relationship. We identified 11 pairs of protein-coding genes and lncRNAs with significantly high Pearson correlation (adj. p < 0.05; Table 1). All pairs were positively correlated, suggesting that some lncRNAs may be acting as enhancers. Six pairs were transcribed on the same strand while five were transcribed on opposite strands.

**Table 1.** Differentially expressed (DE) annotated and predicted lncRNA and protein-coding gene pairs that are within 100kb of each other, and their orientation to each other. Estimated counts were correlated using Pearson correlation and p-values were adjusted using the Benjamini-Hochberg test.

| lncRNA Gene ID | Gene Name | Protein-coding Gene | Direction | Pearson | Pval | Padj |
| --- | --- | --- | --- | --- | --- | --- |
| ENSG00000256894 | NA | BHLHE41 | Sense | 0.880 | 1.50E-46 | 1.80E-45 |
| G25649 | NA | PITX2 | Antisense | 0.842 | 7.38E-39 | 4.43E-38 |
| G10290 | NA | DIO3 | Antisense | 0.837 | 6.03E-38 | 2.41E-37 |
| ENSG00000258498 | DIO3OS | DIO3 | Antisense | 0.790 | 3.69E-31 | 1.11E-30 |
| ENSG00000280798 | LINC00294 | DEPDC7 | Sense | 0.763 | 6.96E-28 | 1.67E-27 |
| ENSG00000240498 | CDKN2B-AS1 | CDKN2B | Antisense | 0.696 | 1.35E-21 | 2.69E-21 |
| G1330 | NA | L1TD1 | Sense | 0.639 | 2.03E-17 | 3.48E-17 |
| ENSG00000256894 | NA | SSPN | Antisense | 0.515 | 7.41E-11 | 1.11E-10 |
| G8996 | NA | OLFM4 | Sense | 0.314 | 0.0002 | 0.000206 |
| ENSG00000280383 | NA | FBLN1 | Sense | 0.233 | 0.0056 | 0.006721 |
| ENSG00000242795 | NA | MECOM | Sense | 0.180 | 0.0332 | 0.036211 |

Along with the well-annotated *ANRIL* and *DIO3OS* lncRNAs discussed above that showed strong correlation with their corresponding antisense protein-coding genes (*CDKN2B* and *DIO3* respectively), several predicted lncRNAs showed strong concordance in expression with a nearby protein-coding gene. For example, we saw a strong positive correlation of *DIO3* with the predicted lncRNA *G10290*, which is ∼10kb downstream of and sense to *DIO3OS*. *G10290* is supported by a CAGE-annotated TSS site in the separate IBD cohort. Similarly, lncRNA *G25649*, which is downstream and transcribed on the opposite strand of the protein-coding gene *PITX2*, was positively correlated with *PITX2*. From this point forward we will refer to this lncRNA as *PANCR-AS1*.

### Co-expression network analysis reveals lncRNAs associated with biological pathways important in IBD

To better understand the potential functional roles of the predicted and annotated lncRNAs in CD, we performed weighted gene co-expression network analysis (WGCNA)[56]. WGCNA generated 52 co-expression modules (Figure 3A; Supplementary Table 4). The size of each module varied from 27 to 3029 coding and non-coding genes. We found that 1752 lncRNAs were members of a co-expressed module, of which 1329 were annotated lncRNAs and 423 were predicted lncRNAs; of these, 48 annotated lncRNAs and 36 predicted lncRNAs were among those differentially expressed in CD.

**Figure 3.**
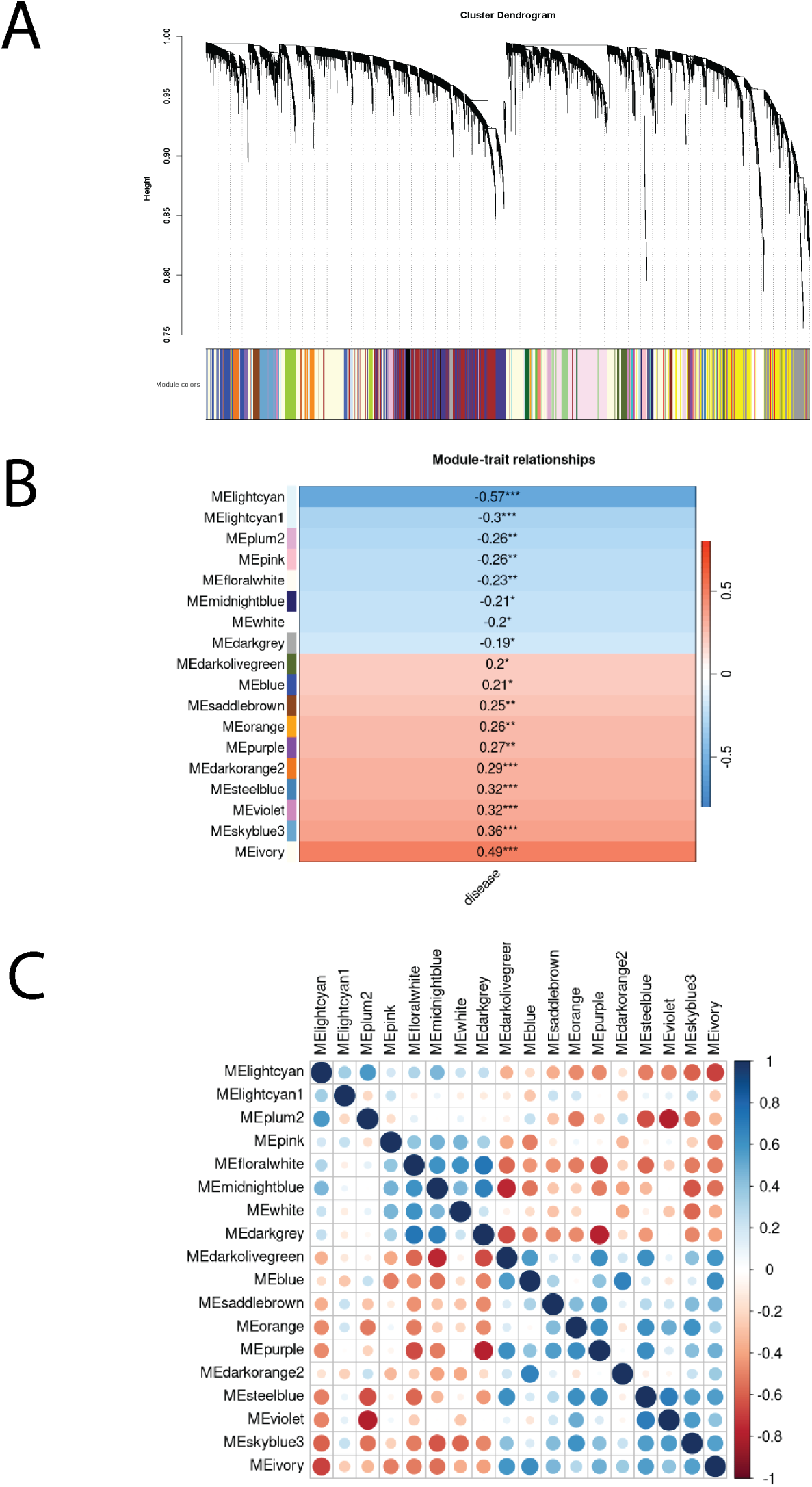
Gene co-expression network analysis of lncRNA and protein coding genes. (A) Cluster dendrogram of WGCNA networks. (B) Clusters whose eignengenes are significantly correlated (p-value < 0.05; Pearson’s) with disease status. (C) Correlations between all pairs of cluster eigengenes for clusters significantly correlated with disease status.

Module eigenvectors, which reflect the general gene expression pattern in a given module, were determined to enable comparison across modules. To determine which modules may be associated with CD, we correlated module eigenvectors with disease status and found 18 that were significantly correlated (p<0.05; Figure 3B). To ensure that module expression between these 18 clusters was sufficiently distinct, we correlated their eigenvectors and found that correlations were all less than 0.72 (Figure 3C). This supports that expression of genes within modules was highly similar but different across modules. To understand what cellular functions each module may represent, we performed pathway enrichment analysis on each of these CD-associated modules. All but three had at least one significant Gene Ontology (GO) term associated with them (p<0.05, p.adjust <0.01; Supplementary Table 5). Using these terms, we labeled each module to reflect their primary associated function (Table 2). We found multiple modules were related to immune system and metabolic processes, as expected.

**Table 2.** Top five GO Terms for significant IBD-related modules (by . IBD-associated clusters with > 20 significant terms, we summarized redundant terms using REVIGO.

| Module Color | GO Terms (Top 5 based on q-value) | Category |
| --- | --- | --- |
| lightcyan | oligosaccharide binding, disruption of cellular component of another organism, antimicrobial humoral response, humoral immune response, antimicrobial humoral immune response mediated by antimicrobial peptide | AMP response |
| lightcyan1 | antigen processing and presentation of peptide antigen via MHC class I, MHC class I protein complex, regulation of adaptive immune response, cellular response to lipopolysaccharide, cell killing | MHC class I activity |
| plum2 | embryonic skeletal system morphogenesis, embryonic skeletal system development, anterior/posterior pattern specification, arachidonic acid epoxigenase activity, arachidonic acid monooxygenase activity | Developmental programs |
| pink | structural constituent of chromatin, nucleosome, protein-DNA complex, protein heterodimerization activity, nucleosome assembly | DNA-protein interactions |
| floralwhite | cell cycle checkpoint signaling, regulation of chromosome organization, chromosomal region, chromosome segregation, DNA replication | Cell replication I |
| midnightblue | endoplasmic reticulum lumen, endoplasmic reticulum protein-containing complex, protein disulfide isomerase activity, intramolecular oxidoreductase activity, transposing S-S bonds | Endoplasmic reticulum activity |
| white | chromosome segregation, chromosomal region, DNA replication, spindle, positive regulation of cell cycle process | Cell replication II |
| darkgrey | preribosome, ncRNA processing, ribosome biogenesis, 90S preribosome, rRNA metabolic process | Ribosome activity |
| blue | collagen-containing extracellular matrix, sarcolemma, muscle system process, basement membrane organization, organ growth | Extracellular matrix activity I |
| saddlebrown | negative regulation of cell activation, antigen processing and presentation, trans-Golgi network, endocytic vesicle, leukocyte proliferation | MHC class II activity |
| purple | NAD+ nucleotidase, cyclic ADP-ribose generating, NAD+ nucleosidase activity, NAD(P)+ nucleosidase activity, phagocytic vesicle, phagocytic vesicle membrane | Energy production and phagocytosis |
| darkorange2 | collagen-containing extracellular matrix, extracellular matrix structural constituent, glycosaminoglycan binding, heparin binding, integrin binding | Extracellular matrix activity II |
| steelblue | late endosome, late endosome membrane, tertiary granule | Neutrophil granules |
| violet | oligosaccharide biosynthetic process, sialyltransferase activity, glycoprotein biosynthetic process, oligosaccharide metabolic process, sialylation | Metabolic processes |
| skyblue3 | apical part of cell, apical plasma membrane, monoatomic anion transport, brush border, cluster of actin-based cell projections | Epithelial barrier |
| darkolivegreen | NA | NA |
| ivory | NA | NA |
| orange | NA | NA |

Of the 1752 lncRNAs found in a module, 243 were within one of the 18 disease-associated modules. Of these, 37 lncRNAs were differentially expressed, and these localized to one of the following nine disease-associated modules: MHC class I activity, MHC class II activity, antimicrobial peptide (AMP) response, developmental programs, epithelial barrier, neutrophil granules, metabolic processes, and two clusters of undetermined function (ivory and orange). We highlight four of these disease-associated modules and their lncRNAs below.

A more detailed review showed that the AMP response module contained 39 members, 35 of which were protein coding genes and four were lncRNAs. Of these, 29 were significantly upregulated in CD, including all four lncRNAs. Two of these were from the annotated lncRNA set (*LINC01232* and *ENSG00000288047*), and two were predicted lncRNAs (*G1330* and *G8996*). Both *G1330* and *G8996* were significantly DE and correlated with the expression of their nearby protein-coding genes (*L1TD1* and *OLFM4* respectively); *L1TD1* activates viral retrotransposition and *OLFM4* promotes mucus secretion, both of which are processes that are aberrantly increased in active IBD [74], [75]. Top gene ontology (GO) biological process (BP) terms implicated numerous DE genes (*DEFA5, DEFA6*, *REG1A, REG1B, REG3A*) with the formation of antimicrobial peptides, an important component of the adaptive immune response to pathogens and are also frequently overexpressed in IBD[76] (Supplementary Figure 5A). To determine what genes were potentially playing a regulatory role on other genes in their module, we looked at the top 25% most well-connected genes, which we will refer to as “hub” genes; in essence, these were the nodes (genes) with the most edges (a strong gene co-expression correlation) to other genes in the module. Among significant protein-coding and lncRNA gene pairs, *L1TD1* and *G1330* were determined to be hub genes (Supplementary Figure 5B).

The Epithelial Barrier module consisted of 361 protein-coding genes and 68 lncRNAs, of which 73 and 12 were significantly downregulated in CD. Four of the differential lncRNAs were predicted (*G21438*, *G17883*, *G14453*, and *G10290*). Top GO Biological Process terms implicated many of these genes in ion transport and metabolic processes (Supplementary Figure 5C), which are critical to maintaining a functional intestinal barrier. Hub genes in this module included *ANRIL* and *DIO3OS*, which have been reported to be downregulated in IBD patients (Supplementary Figure 5D)[77].

The Metabolic Processes module contained 16 downregulated DE protein-coding genes, including genes related to the process of adding glycans to proteins (*ST6GAL2*, *ST6GALNAC6*) (Supplementary Figure 5E). The addition of sialic acid residues (or sialyation) to mucus-secreting proteins (mucins) plays an important role in protecting them from degradation by bacteria, promoting barrier integrity[78], [79]. Loss-of-function mutations in B3GALT5 and ST6GALNAC1 have been discovered in whole-exome sequencing studies of patients with IBD, and further studies of mice lacking the expression of *B3GALT5* and *ST6GALNAC6* (which encodes a protein similar in structure to ST6GALNAC1) decreases the sialylation of MUC2, a mucin implicated in the development of spontaneous colitis upon knockdown[79]. While *B3GALT5* expression did not reach statistical significance (adj. p = 0.12), a lncRNA that is transcribed in the opposite direction of the gene from the same promoter region (*B3GALT5-AS*1) was DE, and they were both considered hub genes in their module (Supplementary Figure 5F).

Lastly, the Developmental Programs module contained seven DE lncRNAs, including five annotated lncRNAs (*ENSG00000267709*, *LINC01819*, *LINC01687*, *LINC01006*, *ENSG00000246090*) and two predicted lncRNAs (*G25649* and *G17790*), all upregulated in IBD. Protein-coding genes included *GPC3,* members of the homeobox (HOX) gene family (*HOXA3*, *HOXB3*, *HOXB4*, *HOXB5*, *HOXB6*, *HOXB8*), and *PITX2*, which are all associated with pattern formation and specification (Figure 4A); *HOXB8* and *PITX2* were DE, and *PANCR-AS1* and *PITX2* were hub genes in their module (Figure 4B). Interestingly, this module was strongly anti-correlated with the post-translational modification module (cor = -0.79); this is in line with previous studies that have shown the hyperactivity of tissue regenerative processes such as epithelial-mesenchymal transition (EMT) in IBD that lead to the development of fibrosis[80], [81], and that sialyation is downregulated during EMT[82].

**Figure 4.**
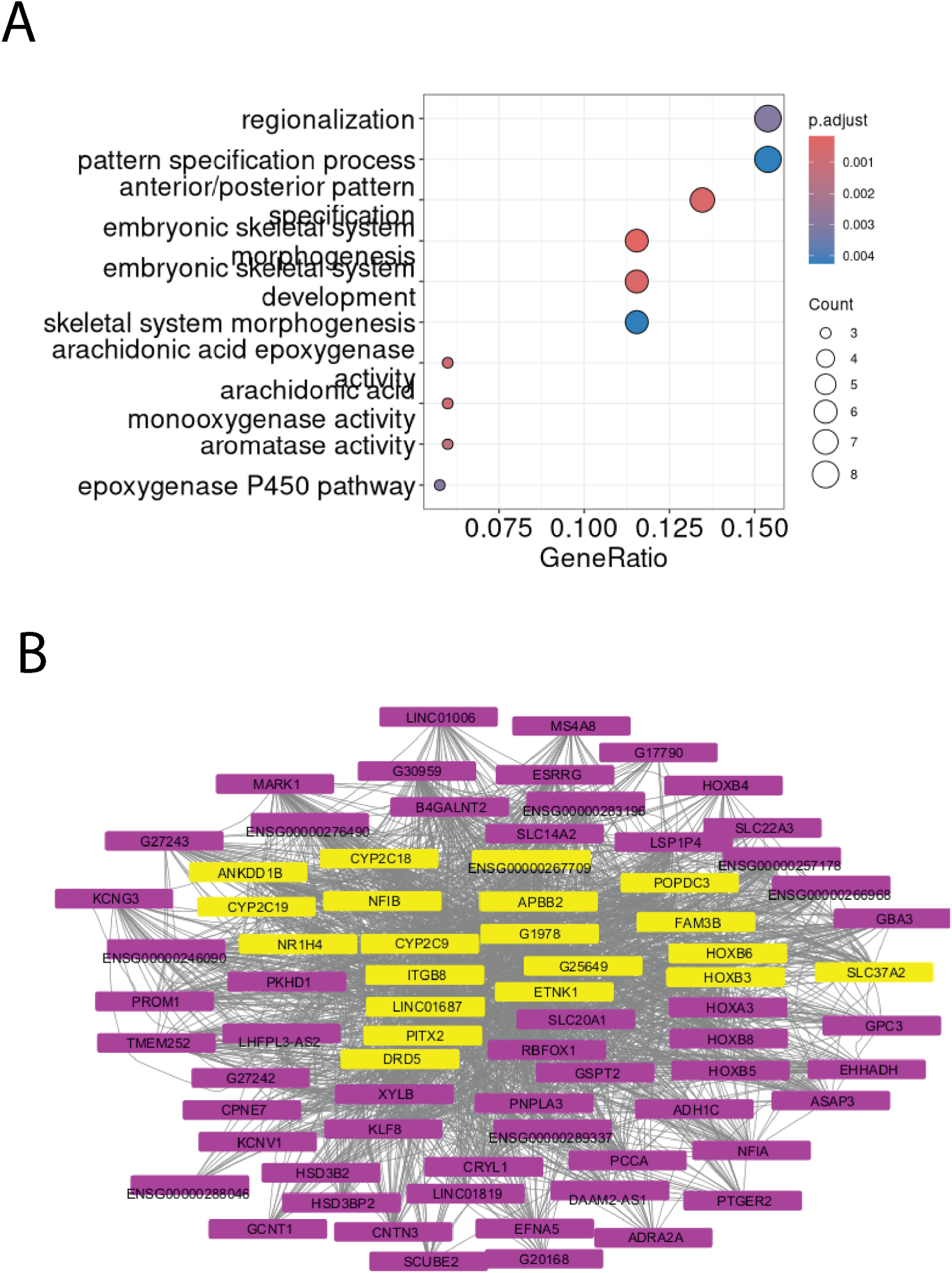
Many significant IBD clusters contain genes associated with processes altered in IBD, with multiple lncRNA-protein coding gene pairs as network hubs. (A) Top GO Biological Processes for the developmental programs (B) Network of genes in the developmental programs module; hub genes, representing the most well-connected genes, are highlighted in yellow.

### *PANCR-AS1* contains evolutionarily conserved regulatory elements with evidence of transcript synteny across multiple species

To hone in on lncRNAs that could play a functional role in on their nearby genes, we accessed evidence of transcript synteny; in essence, this is when a transcript that is found in a different species is flanked by orthologous protein-coding genes, is in the same orientation relative to the orthologous genes, and shows some level of conservation for one or more exons. When looking at conservation between non-coding transcripts flanked by orthologous protein-coding genes in the mouse, 1445 lncRNA transcripts (both annotated and predicted) were predicted to have a noncoding ortholog in the mouse genome (Supplementary Table 6). Two DE predicted lncRNAs in significant modules were considered orthologs of annotated mouse genes, including *PANCR-AS1* (Figure 5A) and *G10290*; while the spliced transcript of *PANCR-AS1* showed lower percent sequence similarity to the mouse ortholog transcript compared to other intergenic transcripts (Figure 5B), the pre-spliced *PANCR-AS1* sequence had greater sequence similarity to the mouse genome compared to others in its category. Interestingly this was true even in comparison to other lncRNAs with known functions that are better evolutionarily conserved, such as those that encode for miRNA host genes or snoRNAs (Figure 5C). *PANCR-*AS1 also had an unusually low insertion/deletion rate relative to other lncRNAs showing a high degree of sequence conservation.

**Figure 5.**
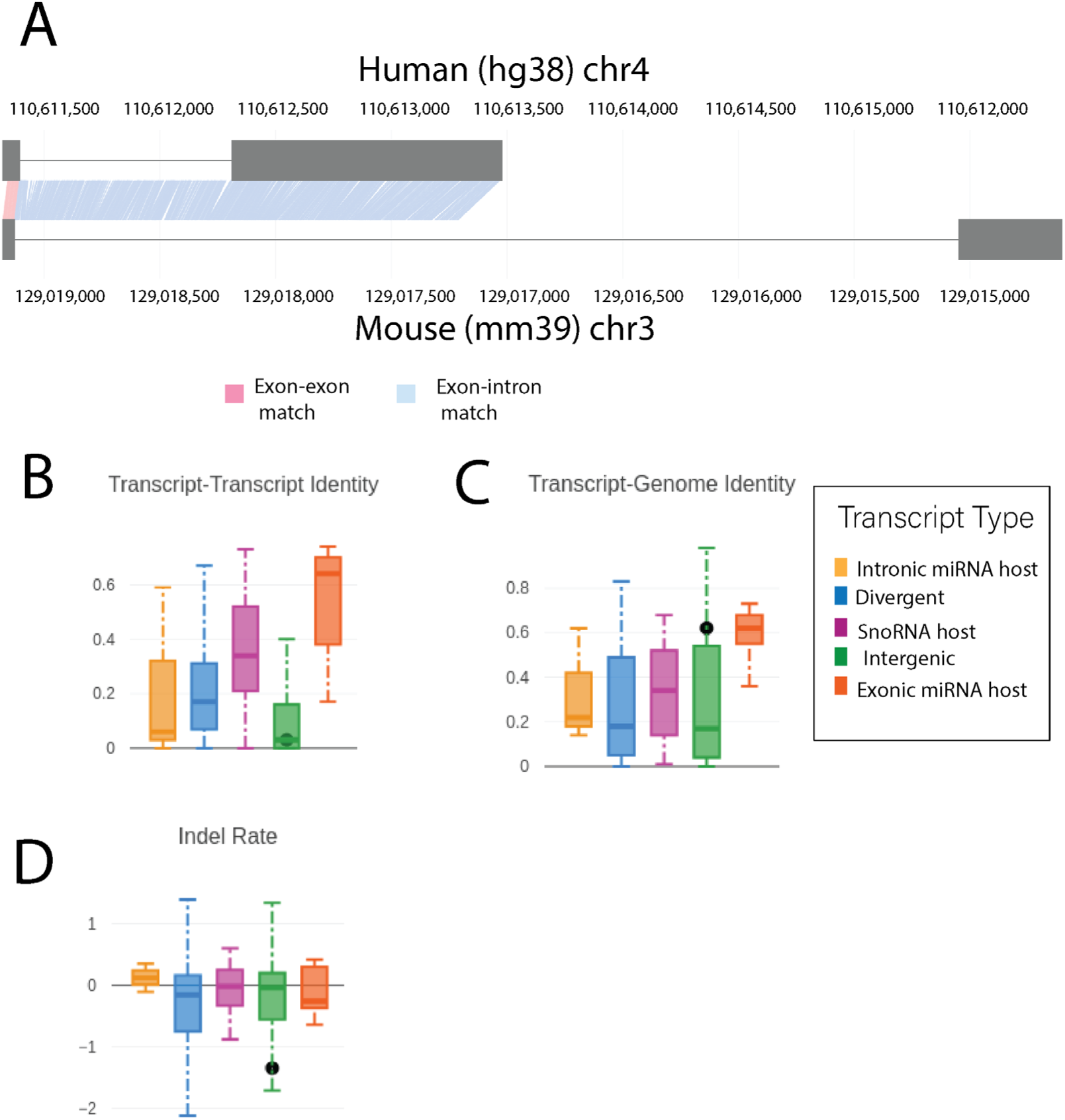
PANCR-AS1 regulatory elements show evidence of conservation. A) Alignment of human PANCR-AS1 to its putative orthologous mouse (mm39) transcript. B) Sequence similarity of predicted human lncRNA transcripts to their putative orthologous mouse transcript. C) Sequence similarity of predicted human lncRNA transcript to the mouse genome. D) Insertion/deletion rate across based on sequence alignments of predicted human lncRNA transcripts to their putative orthologous mouse transcript. In B-D, lncRNAs are broken into five transcript categories based on the functional product: host genes for snoRNAs, host genes for miRNAs where the miRNA is in an intron, host genes for miRNAs where the miRNA is in an exon, lncRNAs transcribed from the same promoter as a protein-coding gene (divergent), and intergenic lncRNAs. PANCR-AS1 is classified as an intergenic lncRNA and is represented by the black dot in all plots.

In fact, when we looked at the overall conservation of *PANCR-AS1* across mammalian species, we found that sequence in the promoter region (+100 bp from the TSS), the donor and acceptor splice sites (+/-2 bp), and the transcript stop site (-2 bp) were highly conserved, as indicated by high average PhastCons scores (0.736 for promoter; 0.996 for donor splice site; 0.770 for acceptor splice site; 0.895 for transcript stop site; Supplementary Figure 6A). The first exon (∼100 bp), promoter, and splice sites are generally well-conserved; specifically, a highly-conserved region of the promoter and the splice site showed general immutability, as evidenced by low tolerance to mutations based on their CADD score, and a contiguous region within the promoter was considered highly constrained on account of their Zoonomia RoCCs score (Supplementary Figure 6B). In addition, it shows matches to transcripts that are annotated in other distantly-related species such as cats, horses, goats, and sheep (Supplementary Figure 7).

### Transcription factor binding patterns suggest that *PANCR-AS1* may function as a tissue-specific enhancer of PITX2 expression

*PANCR-AS1* is robustly expressed across both IBD and non-IBD cohorts (TPM > 1 in 41/140 samples, median 0.42 TPM), while *PANCR* had more sporadic expression (TPM > 1 in 29/140 samples, median 0.17 TPM) and was filtered out prior to differential expression analysis. The transcription start site of *PANCR-AS1* was further supported by CAGE data from FANTOM, as well as PRO-Cap data from the Caco-2 colorectal cancer cell line (Supplementary Figure 8).

We found further evidence of the transcription of *PANCR-AS1* based on the binding patterns of specific transcription factors and chromatin accessibility, whose patterns seemed to be specific to digestive tissues such as the colon. While the *PITX2* promoter was accessible across many tissues, promoter accessibility of *PANCR-AS1* was limited to only a few tissues, including the colonic mucosa (Figure 6A). The ENCODE consensus cis-regulatory element (cCRE) annotation defines the *PANCR-AS1* promoter area as an enhancer. The ChromHMM state annotation of regulatory function labels the promoter region as bivalent (poised for activation) in the colonic mucosa, suggesting the presence of both H3K4me3 (active) and H3K27me3 (repressive) histone modifications (Figure 6B). Furthermore, publicly-available ChIP-seq indicated digestive tissue-specific binding of several transcription factors including members of the NELF complex (NELFA, NELFCD, and NELFE) and SUPT5H/SPT5 (Figure 7A); all of these play a role in regulating transcription elongation[83]. In addition, Pol II binds in digestive tissues (Supplementary Figure 9); given that Pol II is recruited to the site of transcription, it is further evidence that transcriptional processes are being activated in *PANCR-AS1*.

**Figure 6.**
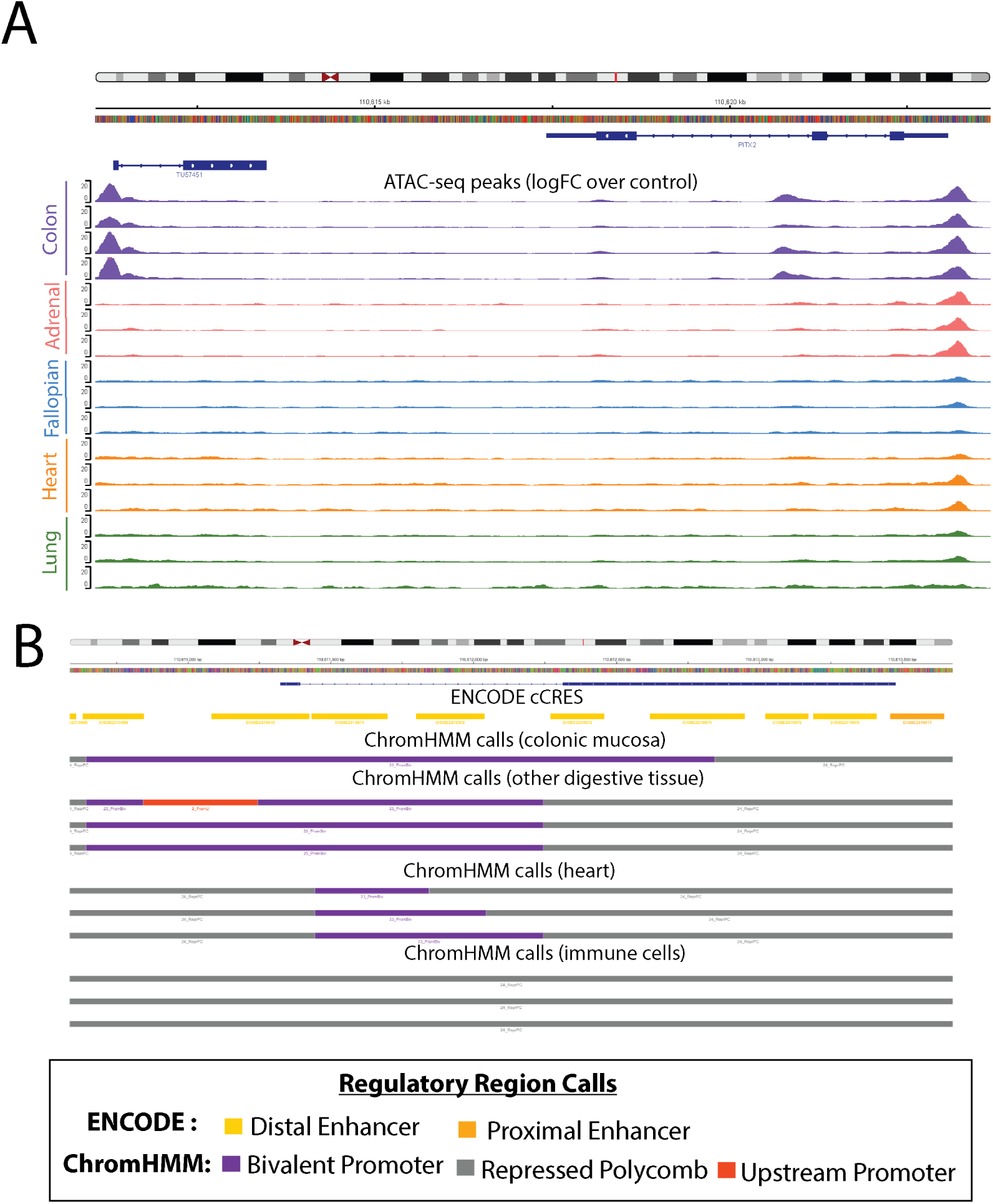
PANCR-AS1 shows tissue-specific chromatin accessibility. A) ATAC-seq peaks (logFC over control) of multiple tissues from the ENCODE project consortium; each color represents samples from a different tissue. B) Overlap of PANCR-AS1 transcript with set of ENCODE cCREs and ChromHMM calls on multiple cell lines and tissues.

**Figure 7.**
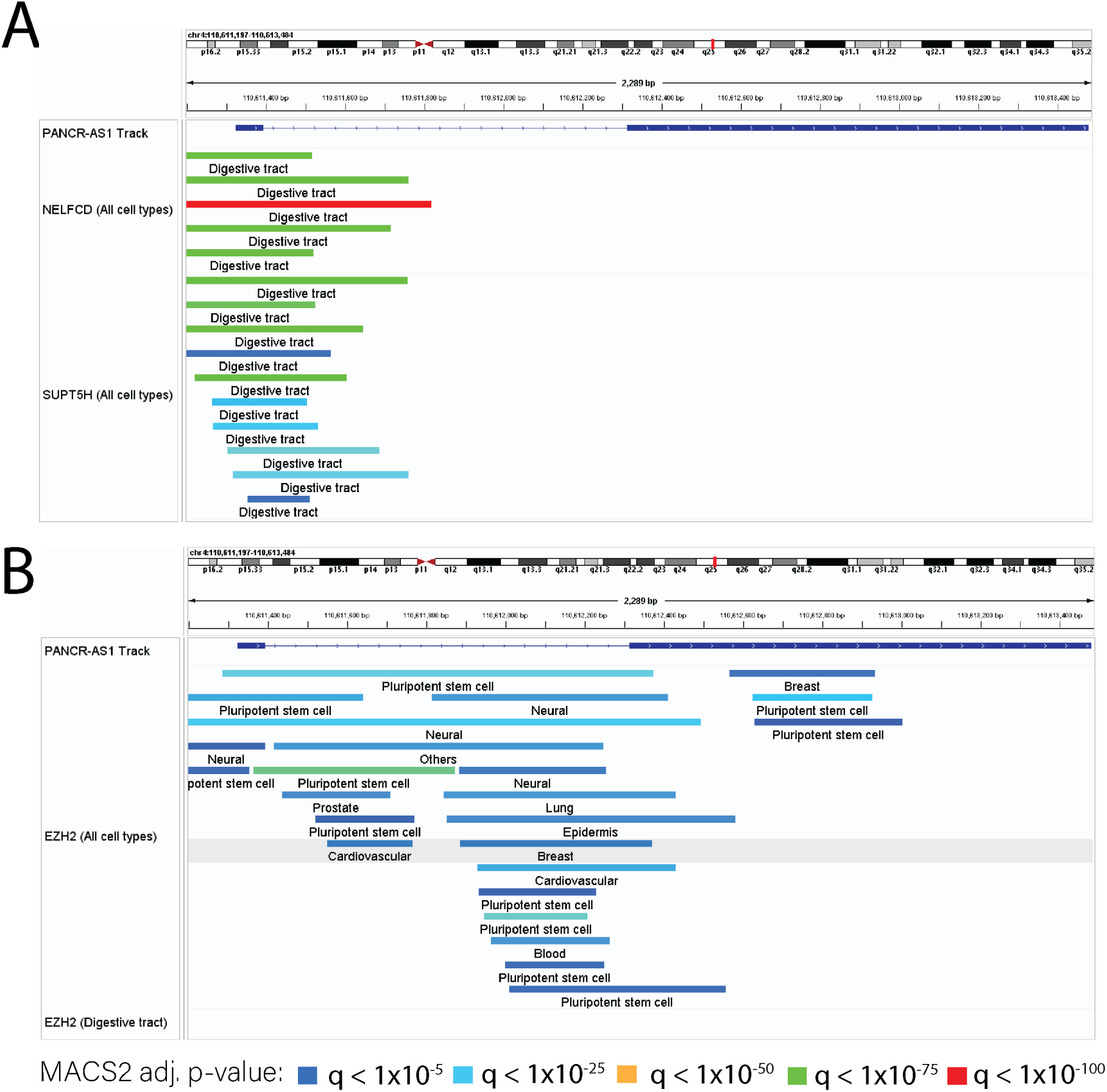
ChIP-seq expression patterns across cell lines and tissues support PANCR-AS1 repression in non-digestive tissues and transcriptional activity in digestive tissues. A) NELFCD and SUPT6H peaks across tissues and cell lines B) EZH2 peaks in non-digestive and digestive tissues. Colors represent the degree of significance of each peak based on MACS2; the colors range from blue to red, where blue represents an adj. p < 1x10^-5^, cyan represents an adj. p < 1x10^-25^, green represents an adj. p < 1x10^-50^, yellow represents an adj. p < 1x10^-75^, and red represents an adj. p < 1x10^-100^.

In contrast, other tissues showed evidence of polycomb repressive chromatin (PRC) marks overlapping part or all of the transcript; in digestive tissues PRC marks only overlapped the second exon (while showing bivalent chromatin marks in the promoter and first exon regions), while some tissues such as the heart showed evidence of bivalent chromatin only in the intron of *PANCR-AS1*, and other tissues (such as immune cells) showed complete repression (Figure 6B). RIP-seq data, which captures RNA interactions with proteins, showed EZH2 in mouse showed interactions between EZH2 and the start of the *PANCR-AS1* transcript (Supplementary Figure 10) in embryonic stem cells, which are in a repressed chromatin state. Furthermore, ChIP-seq data supported the lack of binding of PRC2 components (EZH2, SUZ12) in digestive tissues, whereas they were binding in non-digestive tissues (Figure 7B).

Finally, evidence supports interactions between the *PANCR-AS1* and *PITX2* in the Caco-2 colon cancer cell line. Chromatin Interaction Analysis with Paired-End-Tag sequencing (ChIA-PET) is an assay used to study 3D interactions within the genome by showing where regions of chromatin interact with each other. In the Caco-2 colon cancer cell line, CTCF and POL2RA has peak calls at the TSS site of *PANCR-AS1*, confirming the presence of these transcription factors using another data modality (Figure 8). In addition, one CTCF sample and both POL2RA samples show evidence of loops spanning from the promoter of *PANCR-AS1* to the start site of the transcript encoding the shorter isoform of *PITX2* (PITX2c) (Figure 8). Together these support the hypothesis that *PANCR-AS1* functions as an e-lncRNA that promotes the expression of *PITX2*.

**Figure 8.**
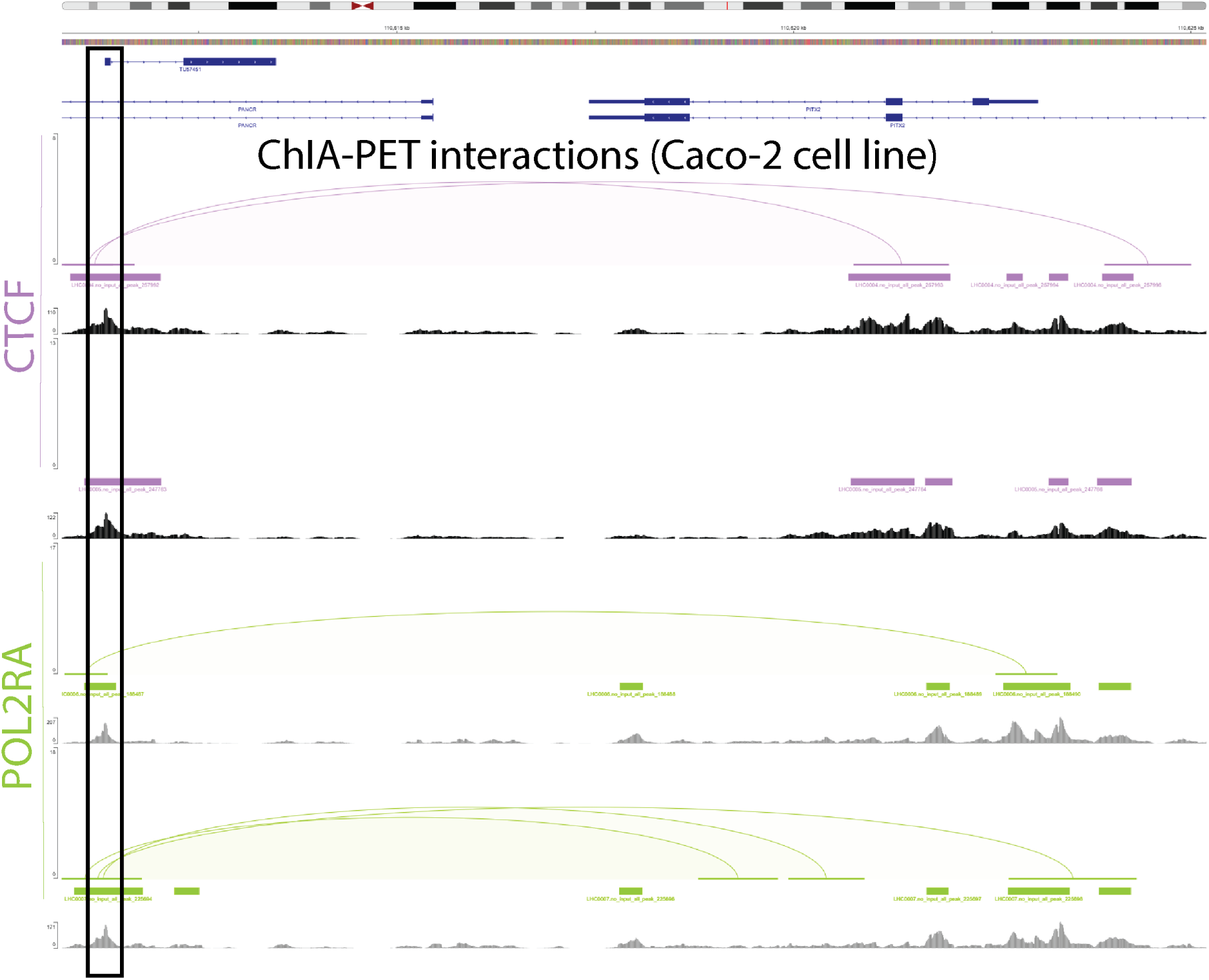
CHIA-PET data from Caco-2 cell line shows interactions between TSS of PITX2 and PANCR-AS1. Plotted above are chromatin loops and peaks of CTCF (purple) and POLR2A (green), as well as the signal of unique reads for CTCF (black) and POLR2A (grey) for pairs of replicates.

## Discussion

In this study, we used a genome-guided assembly approach to predict 2520 lncRNAs not in the GENCODE v39 annotation that were being transcribed in uninflamed colonic mucosa tissue. While previous studies have utilized transcript assembly approaches to annotate lncRNAs and understand their transcriptional differences in the context of IBD[10], [84], our study is notable in two ways: 1) the RNA-seq reads used to create the transcriptome reference in previous studies were from immune cell populations, while we generated sequences from bulk intestinal epithelial cell tissue, and 2) the RNA-seq reads in previous studies were unstranded, preventing them from detecting lncRNAs that overlap with protein-coding genes (such as antisense lncRNAs).

We found 98 lncRNAs that were differentially expressed in CD patients. These include many that had been previously identified, such as *LUCAT1*, *DIO3OS*, and *ANRIL*[7], [11], [12], [77], confirming similar patterns of dysregulation in both inflamed and uninflamed IBD tissues; however the majority of our DE lncRNAs had not been previously associated with CD, including 38 of our predicted lncRNAs. In addition, several lncRNAs previously found to be DE in intestinal immune cell populations after provoking an inflammatory response, including *INFG-AS1* and *IRF-AS1* that upregulate nearby *INFG* and *IRF* protein-coding genes, were not DE in our data. This suggests that these changes may be localized to specific intestinal cell populations. There are likely more lncRNAs that are altered in epithelial cell populations in CD that remain undiscovered, underscoring the need for additional efforts to identify a more comprehensive set in specific bulk tissue.

The functions of even most previously annotated lncRNAs are not well understood. Determining the specific mechanistic roles of lncRNAs will require extensive and dedicated efforts. Our analyses provide evidence for roles of differentially expressed lncRNAs in several cellular processes known to be involved in CD, including those related to the adaptive immune response and the regulation of the intestinal barrier. Given that many lncRNAs function to regulate the transcription of protein-coding genes, these may play critical roles in driving the inappropriate immune response characteristic of CD.

We provided evidence that one of our predicted lncRNAs, *PANCR-AS1*, shows high expression correlation to *PITX2*. *PITX2* transcriptionally activates pathways that remodel the cytoskeleton and regulate cell adhesion to enable gut rotation during development and establish left-right asymmetry, which is crucial for the correct formation of the digestive tract[85], [86]. Unexpectedly, while *PITX2* has been found to be differential in IBD, our study found that it was upregulated in the context of disease, whereas it has been found to be downregulated in other studies[87], [88]. As mentioned above, one reason could be due to our focus on macroscopically uninflamed tissue as opposed to inflamed or previously inflamed tissue used in other efforts. The reason for this change, and how this can influence IBD pathogenesis, is an area for future study.

Finally, we provided extensive evidence that *PANCR-AS1* displays characteristics of e-lncRNAs including: 1) tissue-specific expression at the transcriptional level and regulation at the epigenomic level; 2) high conservation in regulatory regions, as well as evidence for a *PANCR-AS1* ortholog in the mouse and potentially other vertebrates; 3) high expression correlation to nearby *PITX2* expression with chromatin interactions with *PITX2* determined through POL2RA and CTCF profiling; and 4) evidence of tissue-specific activity of transcription factors known to promote transcriptional processes. Given that the splicing of e-lncRNAs has been shown to increase Pol II activity, upregulate enhancer transcription, and promote chromatin accessibility[20], [21], our findings point to a potential mechanism of *PITX2* expression regulation.

One interesting finding is that although enhancers are generally not well conserved across species, those that transcribe spliced e-lncRNAs exhibit higher levels of conservation[22]. Notably, the promoter regions of certain e-lncRNAs have been shown to correspond to enhancer-like elements in more distantly related species, even in the absence of an annotated syntenic transcript[22]. A possible explanation for these findings is that the e-lncRNA has simply not been discovered yet, in part due to low expression in the cell types or tissues that were originally used to generate the transcriptome reference. To fully understand the cell-type and tissue-specific regulatory changes of functional lncRNAs (particularly e-lncRNAs) in disease, it is crucial to supplement standard transcriptome references with transcriptomes assembled from the cell and tissue types of interest, as we have done in this study.

The 50 base pair RNA-seq reads affected our ability to assemble longer full-length transcripts; thus, these predicted annotations are not likely to represent the full lncRNA transcriptome in the colon, especially at the isoform level. Since many lncRNAs modify their isoform usage in different disease contexts, it limits our ability to uncover the full landscape of lncRNA expression changes in IBD. Future long-read sequencing-based studies should enable identification of additional lncRNAs, confirm and improve our predicted transcripts, provide better information on isoforms, and help generate a more comprehensive lncRNA reference. In addition, chromatin accessibility and chromatin interaction data in IBD tissues will provide further confirmation of changes at the epigenomic level in the context of disease and are a promising area for future study. In general, the continued generation of data across multiple modalities will contribute to a better understanding of tissue, cell-type, and context-specific lncRNA expression profiles and their relationship to complex phenotypes and disease.

## Conclusions

In this study, we built a pipeline that used a genome-guided assembly approach on RNA-seq data from a cohort of 90 CD and 50 non-IBD patients to identify lncRNAs that were transcribed in macroscopically uninflamed colonic mucosa tissue. We determined 2520 unique lncRNA sequences representing 1436 distinct genes that were not in the GENCODE v39 gene annotation set. We showed that these predicted lncRNAs shared many similar characteristics with annotated lncRNAs and were supported by multiple types of data. We found 98 lncRNAs that showed differential expression in CD patients compared to the non-IBD controls. Our co-expression analyses provided evidence for roles of differentially expressed lncRNAs in several cellular processes known to be involved in CD, including with the immune system, metabolism, and tissue regeneration. Finally, we provided strong evidence that one of our predicted lncRNAs, *PANCR-AS1*, is functionally relevant and is connected to the regulation of the IBD-associated gene, *PITX2*. This work supports the importance of context-dependent transcript annotation and provides a workflow aided by publicly-available datasets to investigate lncRNAs and identify ones of potential functional relevance.

## Data Availability Statement

Normalized gene count data from this study will be deposited to the Gene Expression Omnibus (GEO), while raw RNA-sequencing data will be uploaded to dbGaP.

We have deposited the Snakemake pipeline for generating the lncRNA reads on GitHub (https://github.com/megara23/lncrna-pipeline/), as well as the scripts used for data analysis and for generating figures and tables. (https://github.com/megara23/lncRNAProjectManuscript2025/).

## Acknowledgements

We gratefully acknowledge the technical support from the UNC High Throughput Sequencing Facility. Histological services were provided by the Histology Research Core Facility in the Department of Cell Biology and Physiology at UNC.

## Author Contribution Statement

T.S.F. and S.Z.S. conceptualized the study. B.H., C.B., G. Lau, G. Lian, and D.W. assisted in sample acquisition. M.M.K.N., S.S., N.C.N., A.A., and M.R.S. contributed to data generation and analysis. M.M.K.N. and T.S.F. provided data interpretation and prepared the manuscript.

## Abbreviations

(CD): Crohn’s disease
(IBD): inflammatory bowel disease
(lncRNAs): long non-coding RNAs
(GWAS): genome-wide association studies
(e-lncRNAs): enhancer lncRNAs
(TPM): transcript per million
(CADD): combined annotation dependent depletion
(RoCCs): runs of contiguous phyloP constraint
(PRO-cap): precision run-on and capped RNA sequencing
(RIP-seq): RNA immunoprecipitation sequencing
chromatin immunoprecipitation sequencing: (ChIP-seq)
(ChIA-PET): chromatin interaction analysis by paired-end tag sequencing
(TSS): transcription start sites
(CAGE): cap analysis of gene expression
(WGCNA): weighted gene co-expression network analysis
(PCA): principal component analysis
(DE): differentially expressed
(GO): gene ontology

