## Supplementary Figures for "Reference-guided Genome Assembly of Long Non-coding RNA Transcripts Reveals Target Genes Associated With Crohn’s Disease"

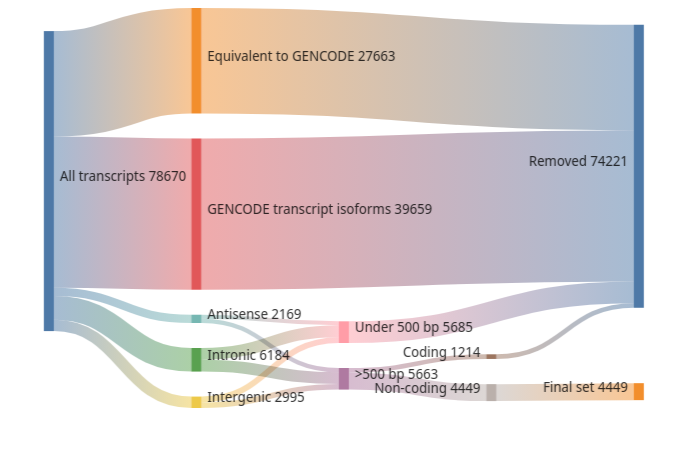

**i	ii	iii**

***Supplementary Figure 1.*** *The number of assembled transcripts filtered out at each stage based on their (i) similarity to GENCODE v39 transcripts (ii) transcript length (iii) coding potential.*

A	B

***Supplementary Figure 2.*** *Overlap of lncRNA transcripts (A) and genes (B) that are still considered novel relative to the GENCODE v47, Lncipedia 5.2, and FANTOM (robust set) transcript databases.*

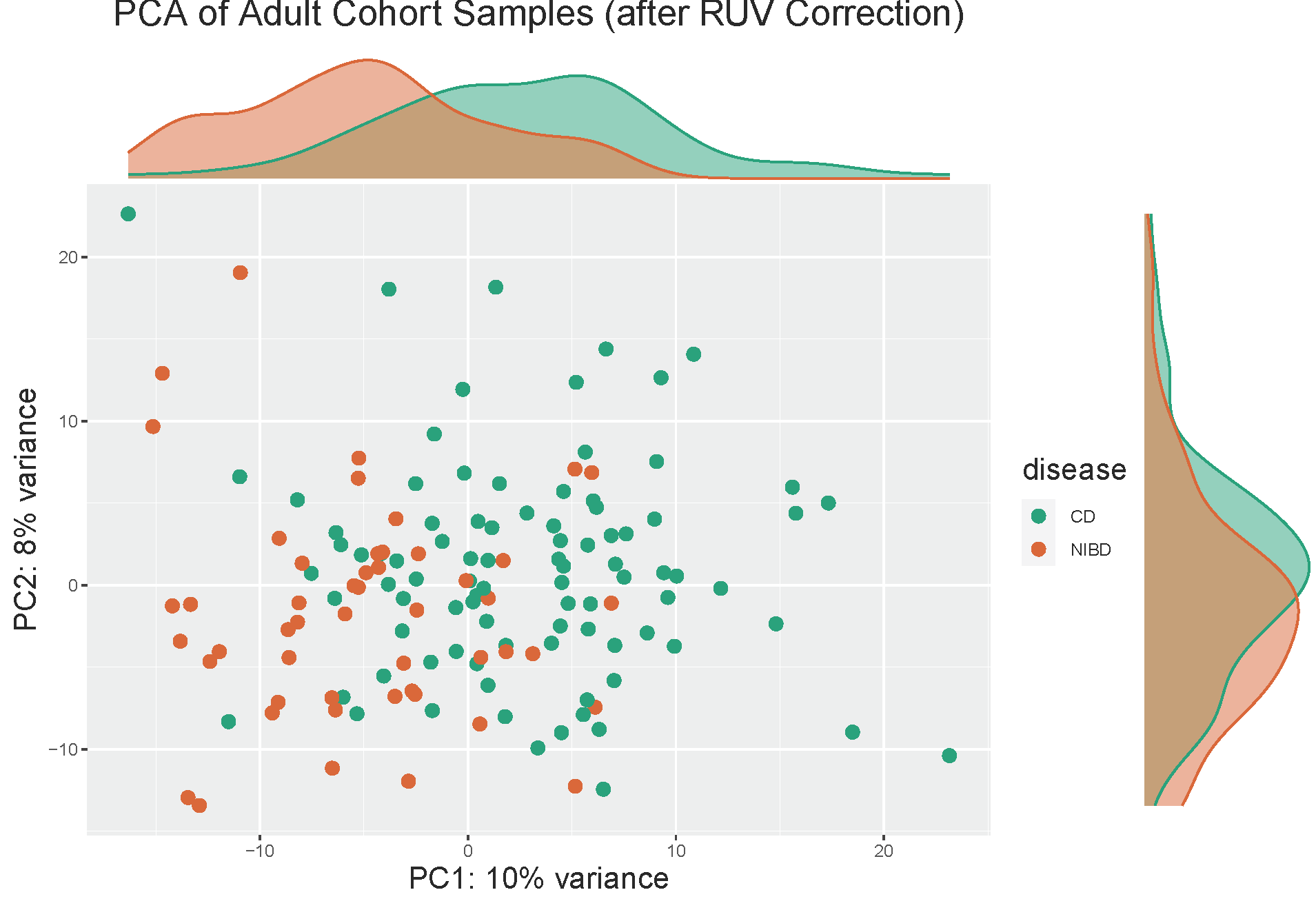

***Supplementary Figure 3.*** *PCA of the top 250 most variable protein-coding genes.*

**A	B**

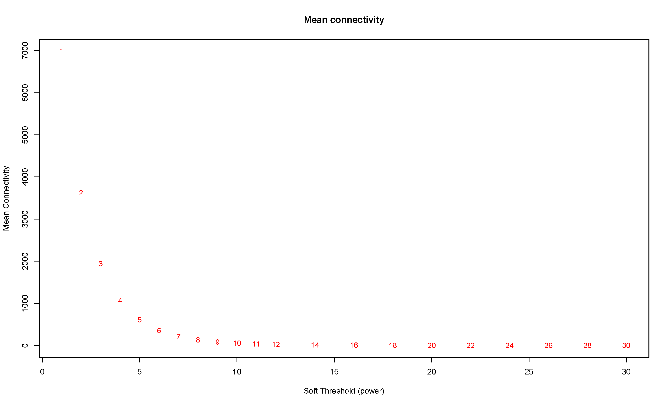

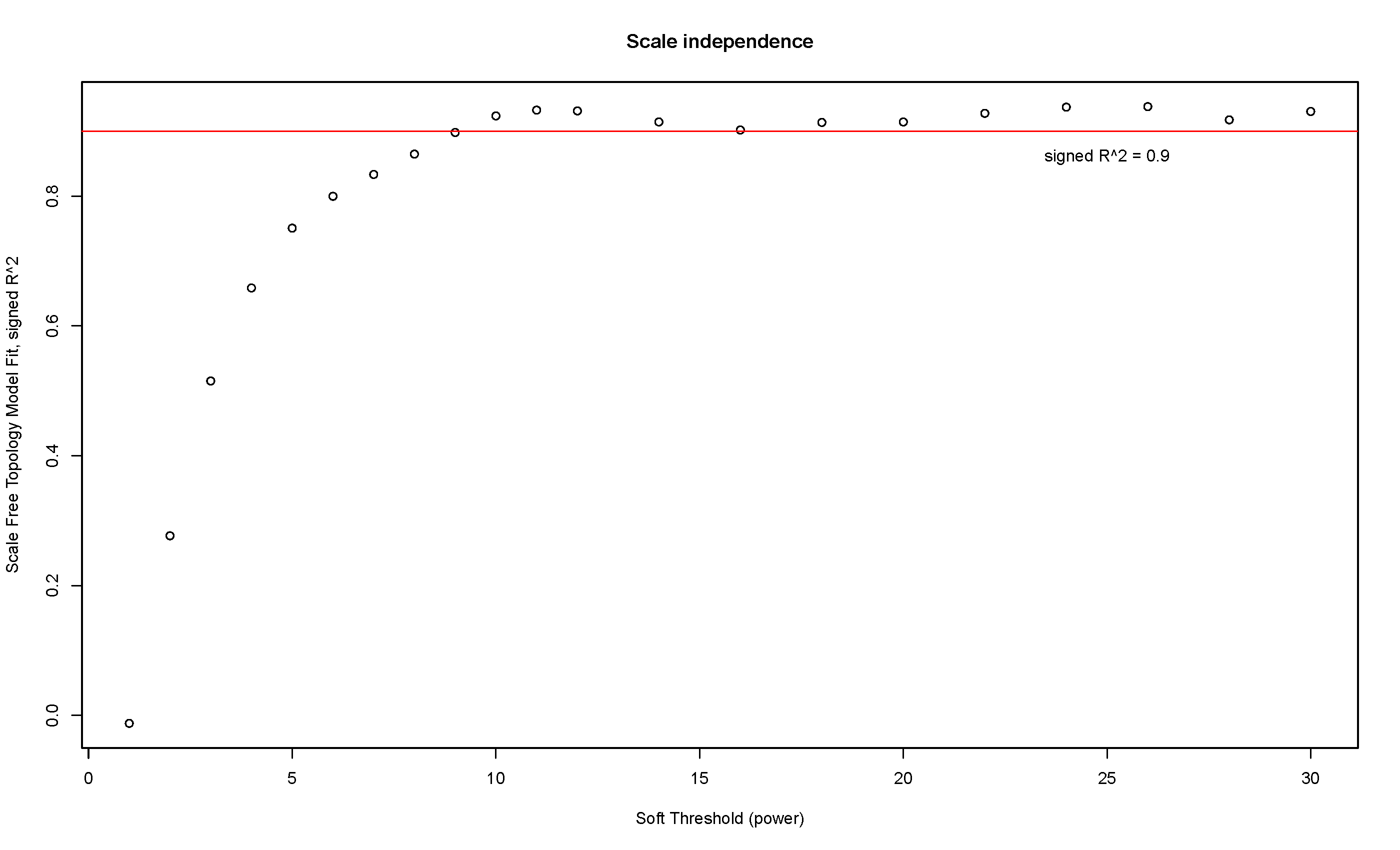

**C**

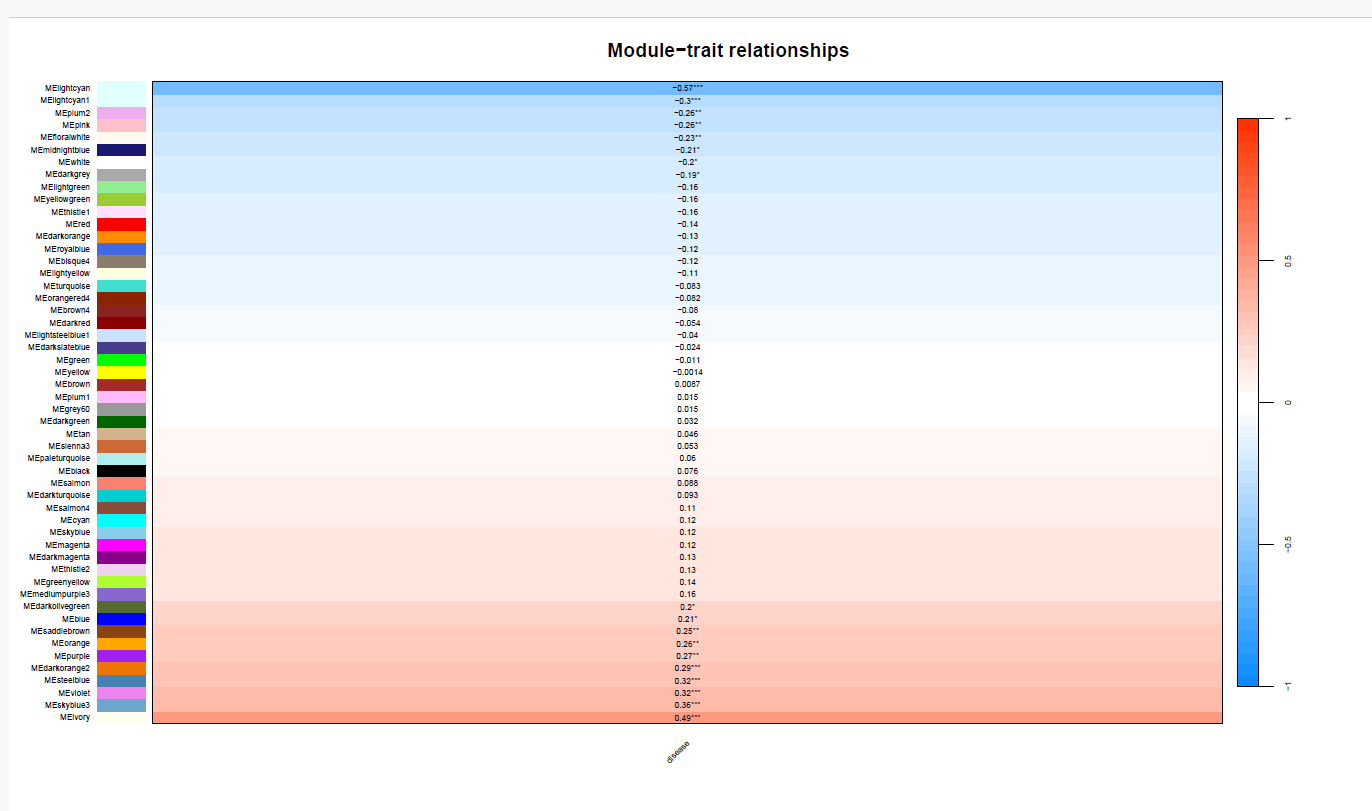

**D**

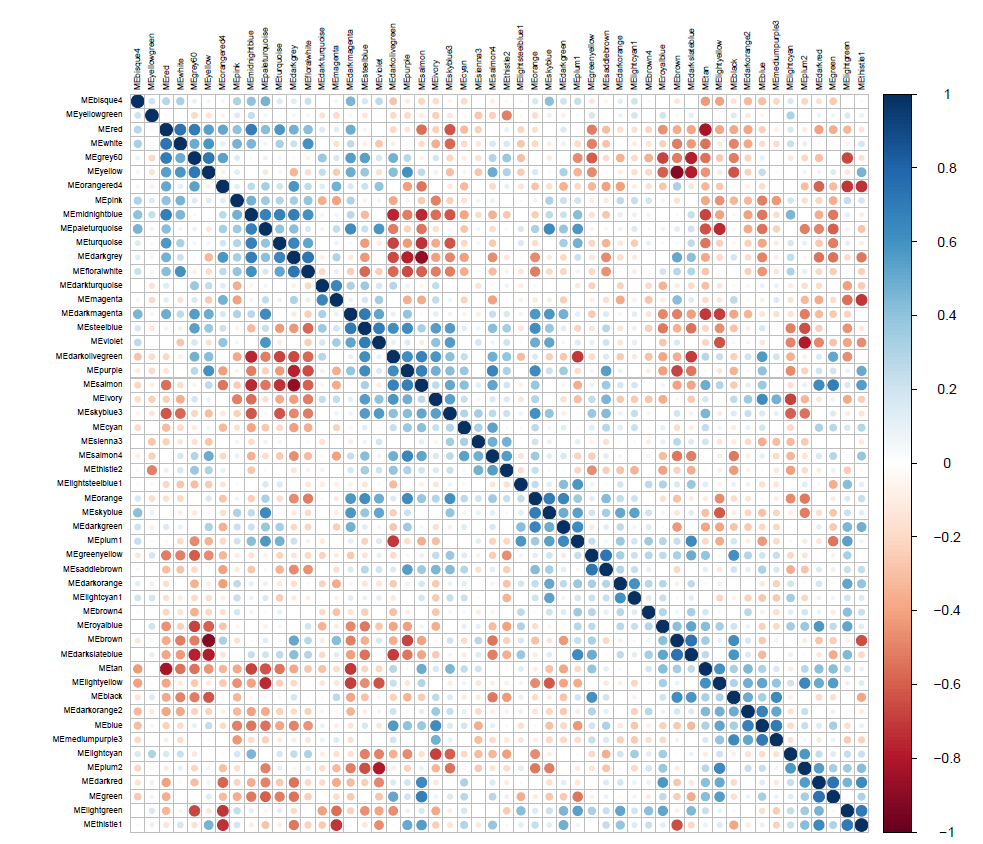

***Supplementary Figure 4.*** *Setting the soft threshold for WGCNA threshold based on (A) topology and (B) the mean connectivity. (C) Correlating cluster eigengene expression with disease status for each cluster. (D) Eigengene-eigengene correlations amongst all clusters.*

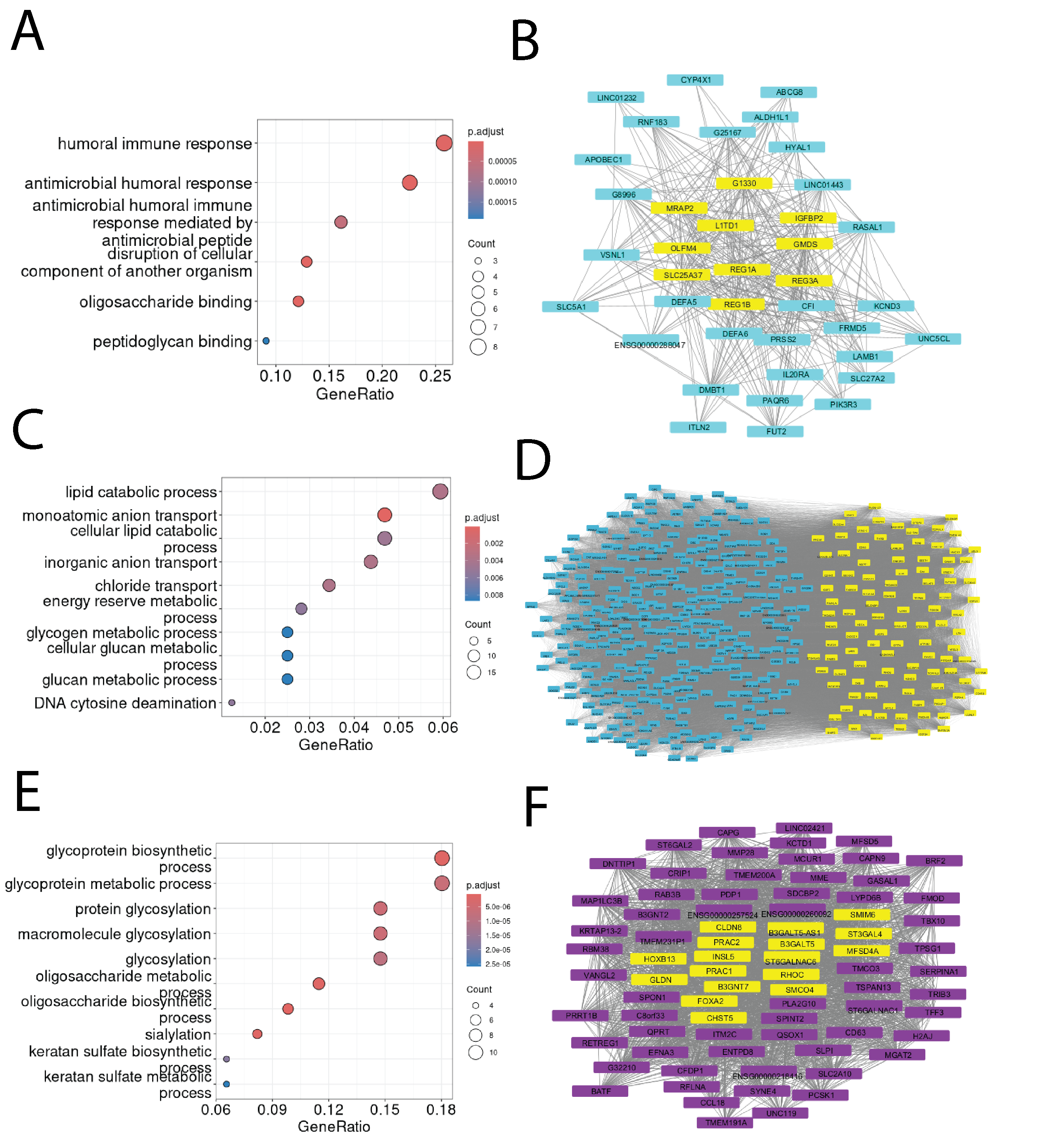

***Supplementary Figure 5.*** ***Other significant IBD clusters with genes associated with processes altered in IBD.*** *Left: Top GO Biological Processes for the AMP immune response (A), epithelial barrier (C), and metabolic processes (E) modules. Right: Network connectivity between genes within the AMP immune response (B), epithelial barrier (D), and metabolic processes (F) modules; the most well-connected genes, or “hub” genes, are highlighted in yellow.*

**A**

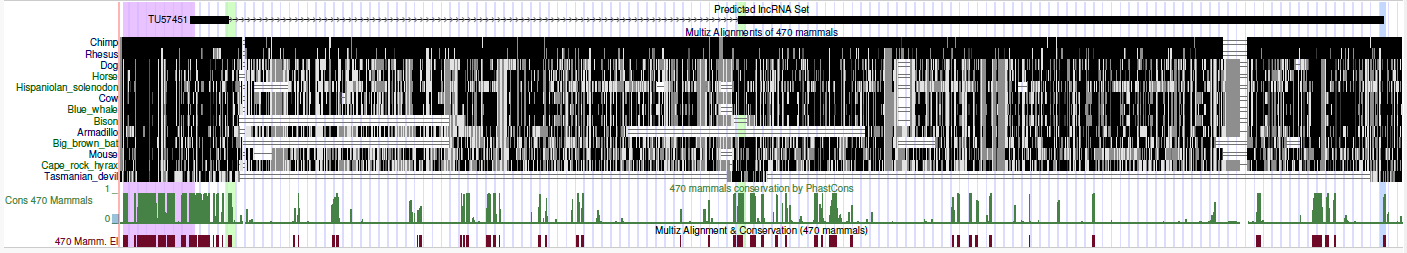

**B**

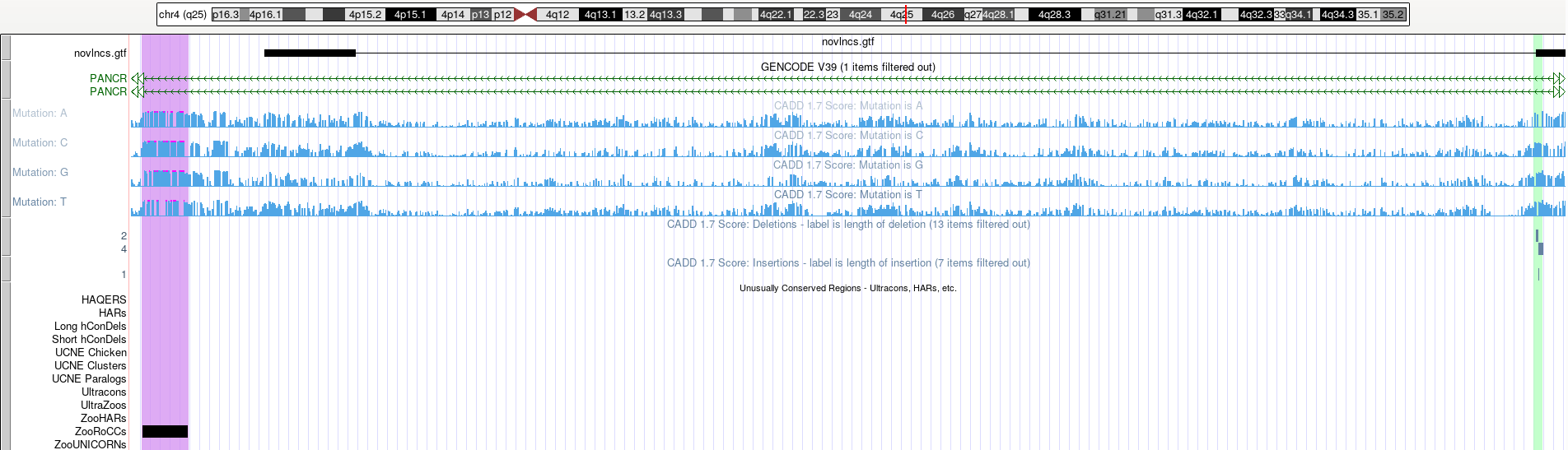

***Supplementary Figure 6.*** *UCSC browser of PANCR-AS1 evolutionary conservation in regulatory regions, with highlights in the promoter (purple), splice sites (light green), and transcript stop site (light blue) regions. (A) Below the transcript is the alignment of the transcript across a select number of species, the PhastCons conservation score (in green), and conservation blocks called by PhastCons (in brown) (B) The promoter region of the transcript has a contiguous series of highly-conserved bases (black box, at bottom) that contains multiple bases with high CADD scores (>20), suggesting their functional importance.*

**A**

**B**

**C**

**D**

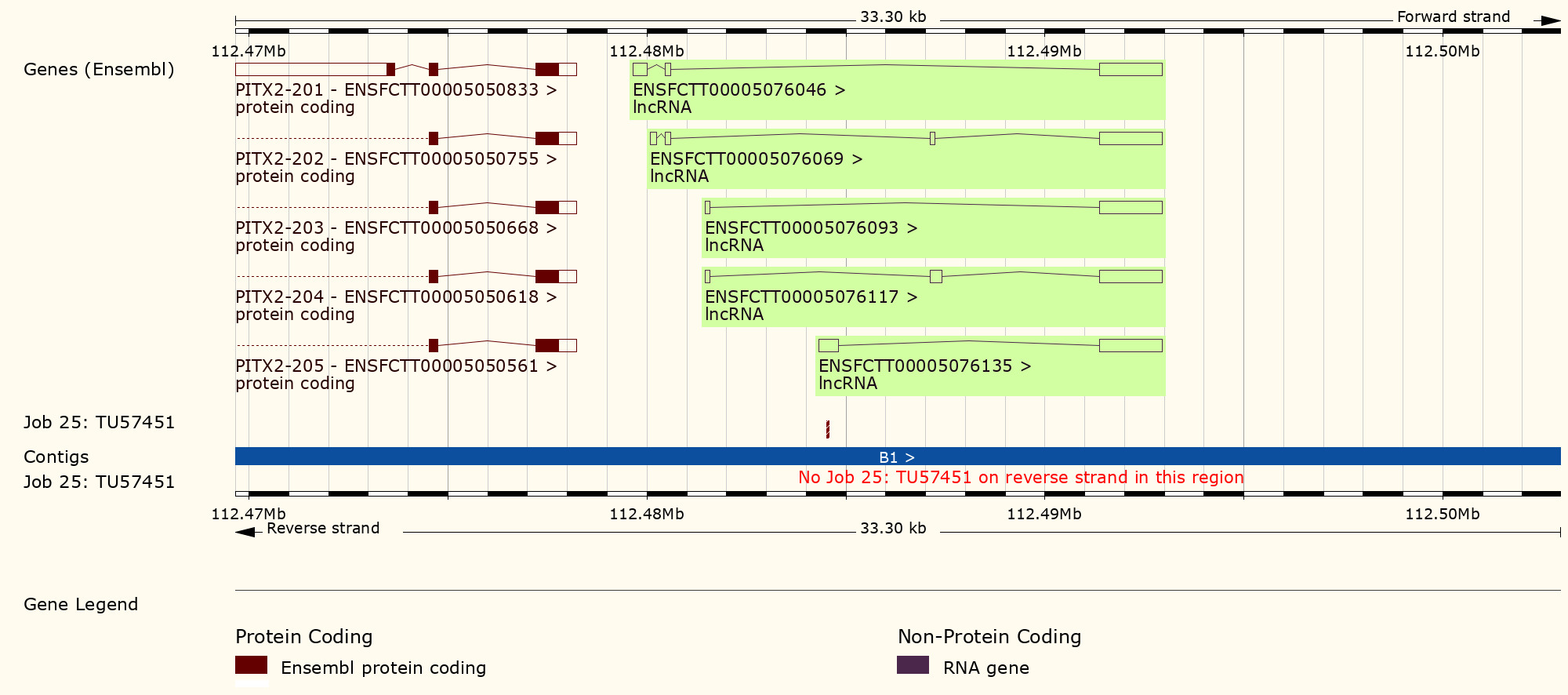

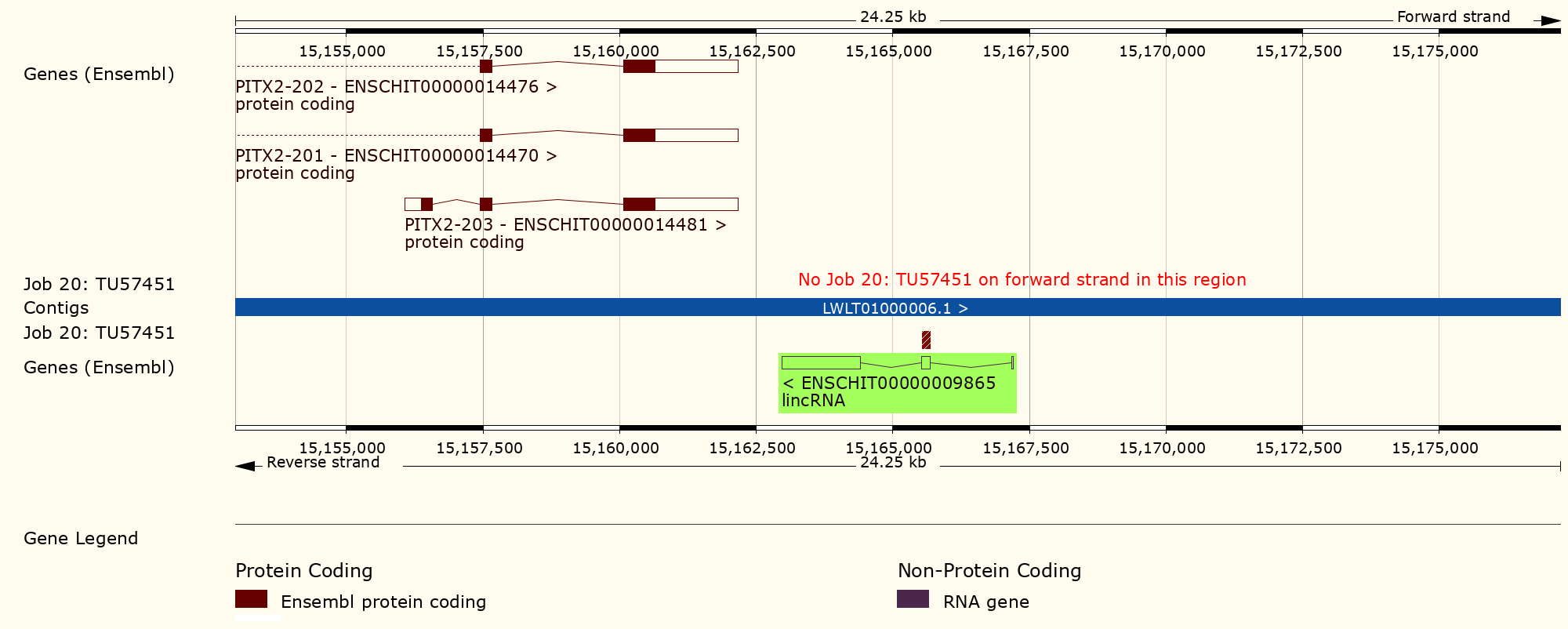

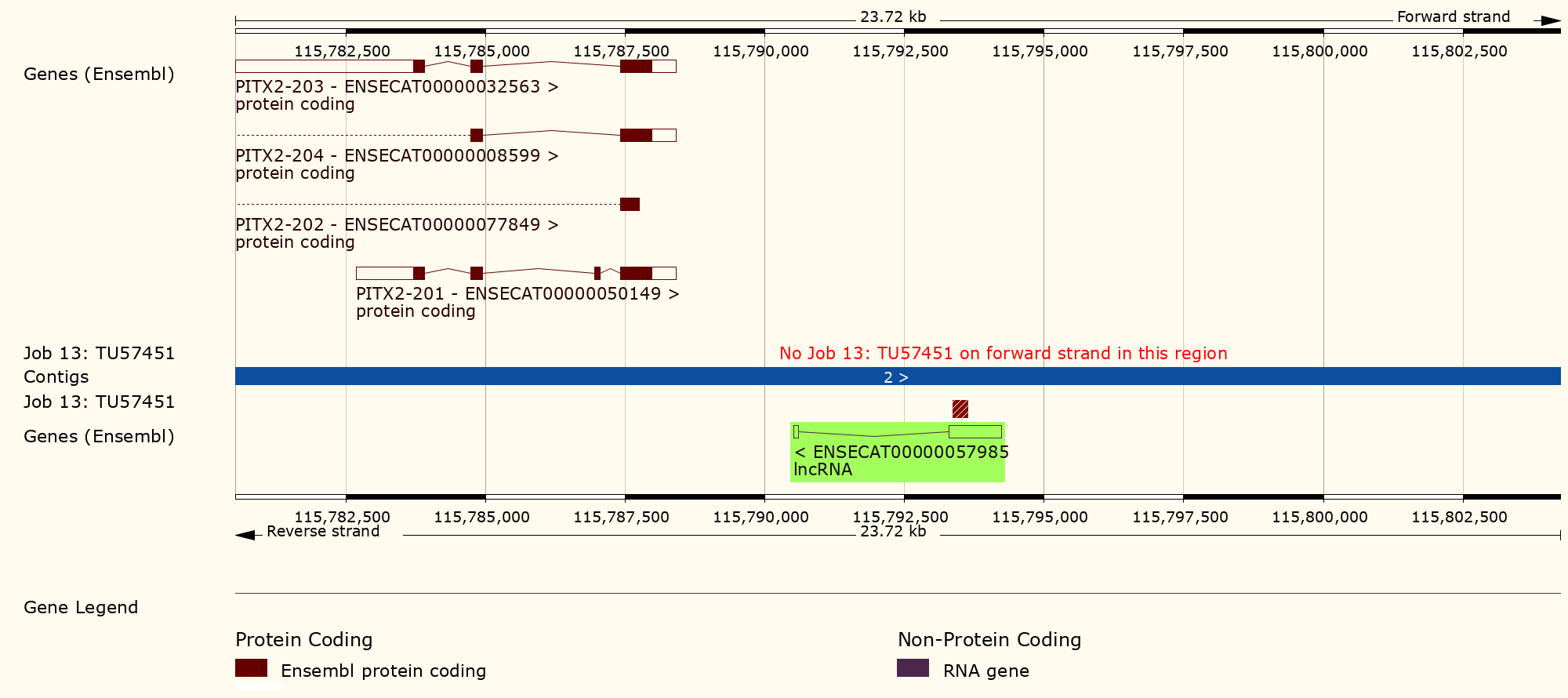

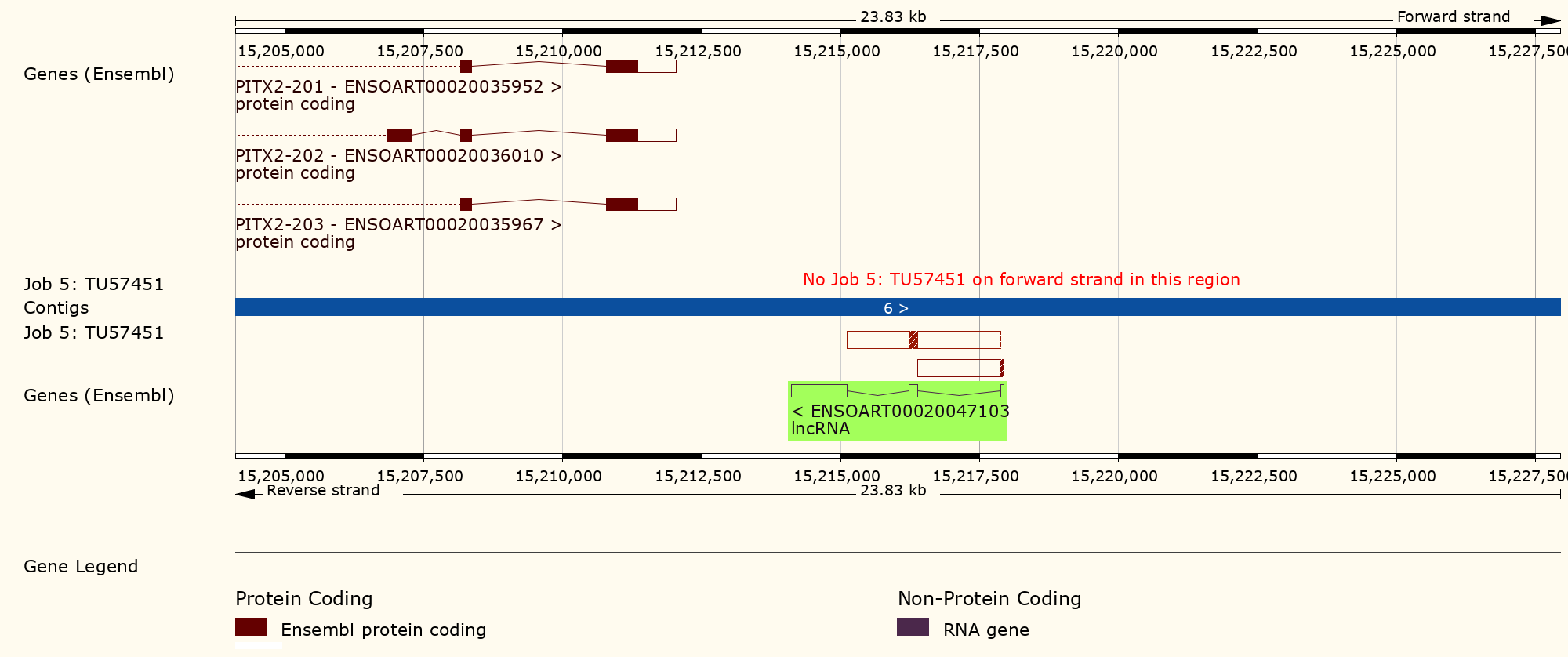

***Supplementary Figure 7.*** *PANCR-AS1 also partially aligns to one or more exons of annotated transcripts in (A) domestic cat (B) goat (C) horse (D) sheep. Transcripts are highlighted in green, with red blocks representing exon-exon sequence alignments. Alignments were made using the Ensembl BLAST/BLAT search tool, with the following parameters changed: “Search against: Ensembl Non-coding RNA Gene Database”, and “Search Sensitivity: Distant Homologies”.*

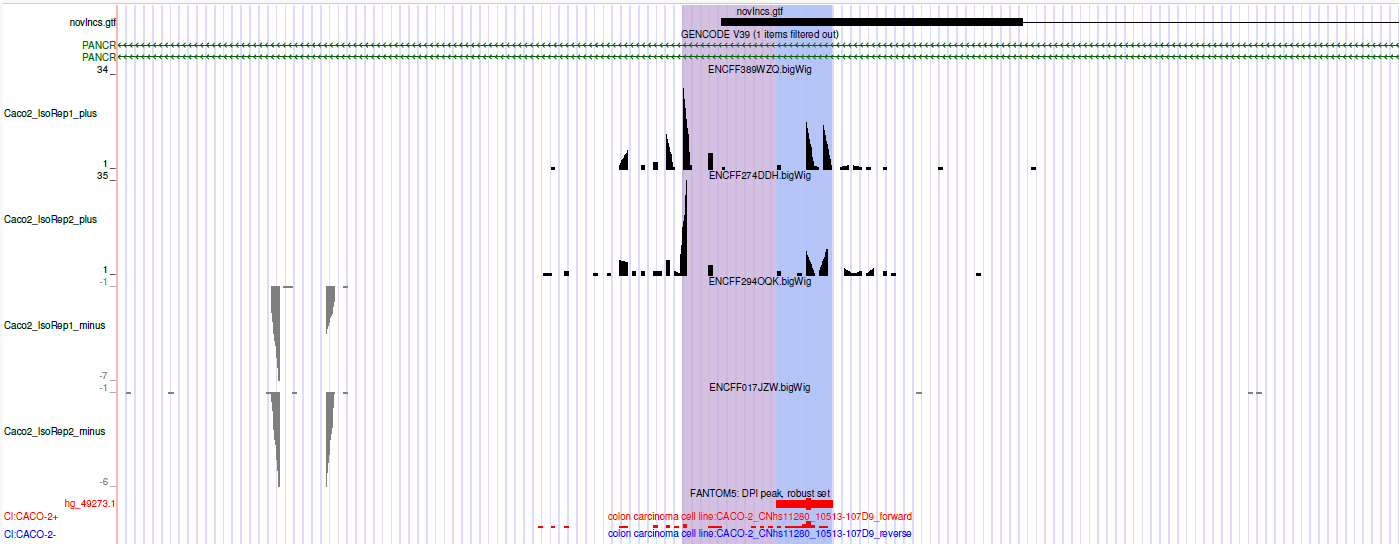

***Supplementary Figure 8.*** *Evidence of nascent transcription of PANCR-AS1 in Caco-2 cell lines, as shown by PRO-cap (purple) and FANTOM TSS expression (blue).*

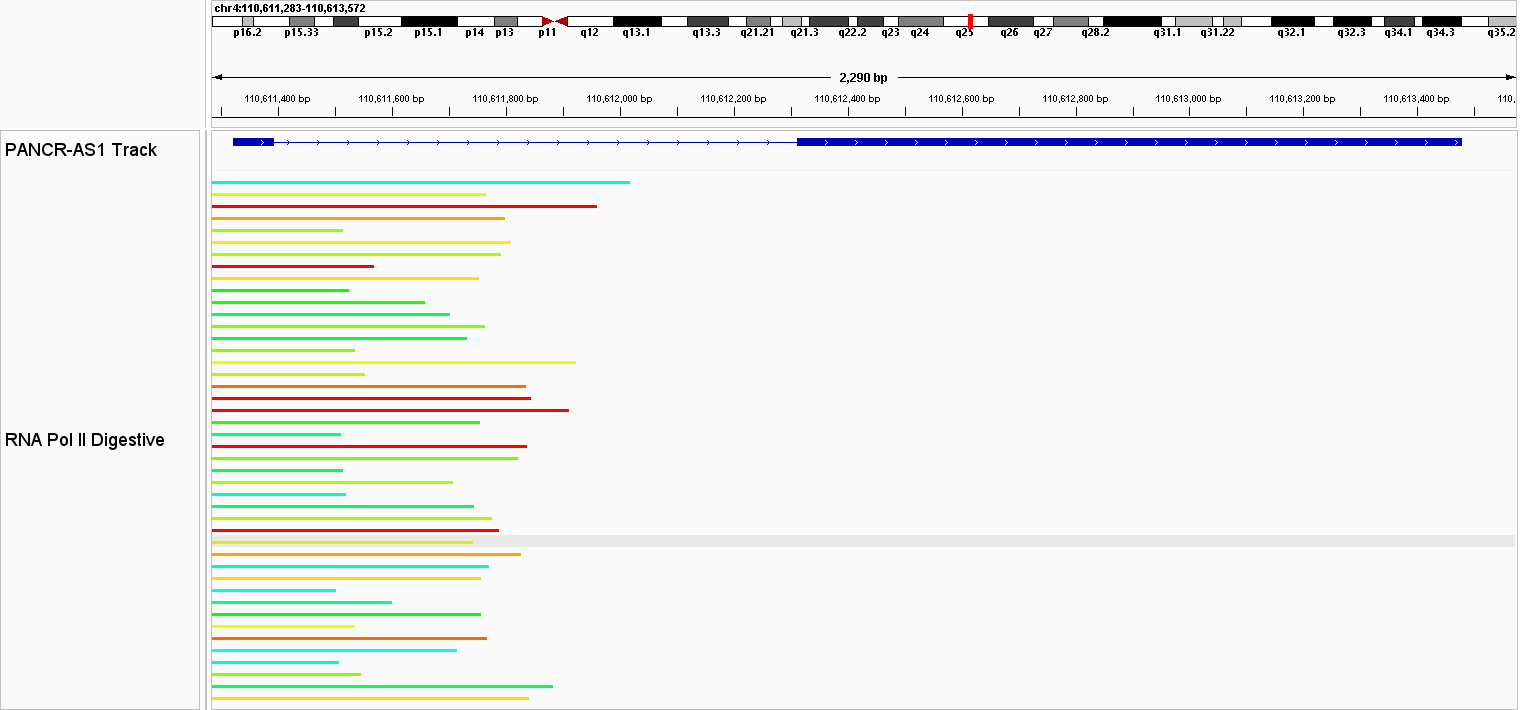

***Supplementary Figure 9*.** *Pol II peaks across digestive tissues and cell lines. Colors represent the degree of significance of each peak based on MACS2; the colors range from blue to red, where blue represents an adj. p < 1x10-5, cyan represents an adj. p < 1x10-25, green represents an adj. p < 1x10-50, yellow represents an adj. p < 1x10-75, and red represents an adj. p < 1x10-100.*

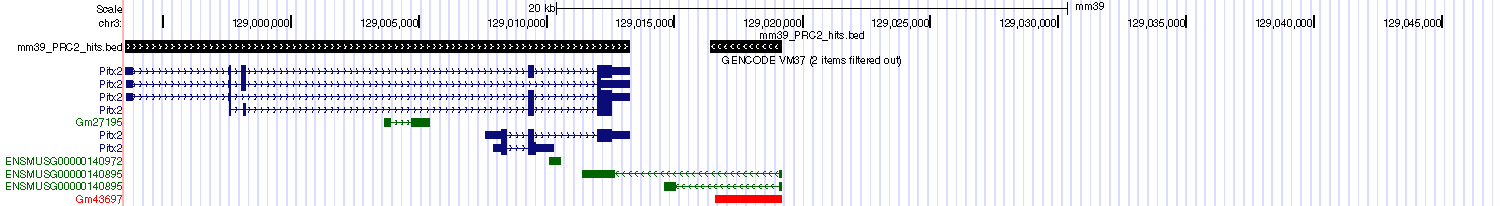

***Supplementary Figure 10.*** *RIP-seq alignments (black) show EZH2 binding to both PITX2 (blue) and PANCR-AS1 (red) in mouse embryonic stem cells using publicly-available data (Zhao et al., 2010). RIP-seq coordinates were lifted over from mm9 to mm10, then mm10 to mm39.*
